# Positive effects of crop diversity on productivity driven by changes in soil microbial composition

**DOI:** 10.1101/2020.11.27.401224

**Authors:** Laura Stefan, Martin Hartmann, Nadine Engbersen, Johan Six, Christian Schöb

**Affiliations:** Institute of Agricultural Sciences, Department of Environmental Systems Science, ETH Zurich, Zurich, Switzerland

**Keywords:** intercropping, soil microbial communities, biodiversity–productivity relationship, sustainable agriculture, annual crop yield

## Abstract

Intensive agriculture has major negative impacts on ecosystem diversity and functioning, including that of soils. The associated reduction of soil biodiversity and essential soil functions, such as nutrient cycling, can restrict plant growth and crop yield. By increasing plant diversity in agricultural systems, intercropping could be a promising way to foster soil microbial diversity and functioning. However, plant–microbe interactions and the extent to which they influence crop yield under field conditions are still poorly understood. In this study, we performed an extensive intercropping experiment using eight crop species and 40 different crop mixtures to investigate how crop diversity affects soil microbial diversity and functions, and whether these changes subsequently affect crop yield. Experiments were carried out in mesocosms under natural conditions in Switzerland and in Spain, two countries with drastically different soils and climate, and our crop communities included either one, two or four species. We sampled and sequenced soil microbial DNA to assess soil microbial diversity, and measured soil basal respiration as a proxy for soil activity. Results indicate that in Switzerland, increasing crop diversity led to shifts in soil microbial community composition, and in particular to an increase of several plant-growth promoting microbes, such as members of the bacterial phylum *Actinobacteria*. These shifts in community composition subsequently led to a 15 and 35% increase in crop yield in 2 and 4-species mixtures, respectively. This suggests that the positive effects of crop diversity on crop productivity can partially be explained by changes in soil microbial composition. However, the effects of crop diversity on soil microbes were relatively small compared to the effects of abiotic factors such as fertilization (3 times larger) or soil moisture (3 times larger). Furthermore, these processes were context-dependent: in Spain, where soil resources were limited, soil microbial communities did not respond to crop diversity, and their effect on crop yield was less strong. This research highlights the potential beneficial role of soil microbial communities in intercropping systems, while also reflecting on the relative importance of crop diversity compared to abiotic drivers of microbiomes, thereby emphasizing the context-dependence of crop–microbe relationships.

## 1. Introduction

The past century has seen the emergence and the development of modern, intensive agriculture that was accompanied by a strong increase in productivity per unit land area (Barrios, 2007). However, the negative effects associated with this intensification now call for a transformation towards more sustainable agricultural production (Tilman *et al*., 2011).

Current intensive agriculture indeed has negative effects on biodiversity (Newbold *et al*., 2015) and ecosystem functioning (Barrios, 2007), including that of soils (Gardi *et al*., 2013). In particular, chemical inputs and the loss of crop diversity associated with intensive agriculture are known to negatively affect soil biodiversity (Cavicchioli *et al*., 2019), with consequences on soil ecosystem functioning, such as nutrient cycling, carbon storage, soil structure regulation, and pest and disease control (Wall & Bardgett, 2012; Wagg *et al*., 2014; Fanin *et al*., 2017). Losses of these essential soil functions can restrict plant growth and subsequently crop yield (Giller, 2001; Bardgett & Van Der Putten, 2014).

Numerous studies demonstrated a strong link between soil microbes and plants (Wardle *et al*., 2004; Fry *et al*., 2018). First, direct specific linkages between a plant species and a particular type of microbe, such as symbiotic interactions, might occur (Wubs & Bezemer, 2016). Moreover, plants have the ability to alter the soil chemical and physical conditions (López-Angulo *et al*., 2020). Some of these changes can be due to above-ground mechanisms, such as litter inputs or variations in microclimate associated with plant canopy cover (Trinder *et al*., 2009; Maestre *et al*., 2009; Delgado-Baquerizo *et al*., 2018). Others may be the result of below-ground processes, such as increases in soil loosening and aeration due to root growth, release of root exudates or plant signaling molecules, and changes in ion uptake (Doornbos *et al*., 2012; Eisenhauer *et al*., 2017). Therefore, a greater richness of plants, and subsequently, roots, would lead to a greater diversity of microhabitats, particularly in the rhizosphere (Philippot *et al*., 2013). The resulting microhabitat heterogeneity would then further enhance microbial diversity (Wardle, 2006; Eisenhauer *et al*., 2010), which could also increase soil functioning, following the diversity-ecosystem functioning relationship (Tilman *et al*., 2006). For instance, Steinauer (2015) showed that increasing plant diversity led to a significant increase in soil microbial biomass and enzymatic activity. Similarly, results from a grassland biodiversity experiment demonstrated that higher plant diversity resulted in increased microbial activity and carbon storage (Lange *et al*., 2015). Most of these studies were performed in natural grasslands; however, the extent to which the same effects would arise in diversified cropping systems is still unclear. Indeed, intercropping, which is defined as growing at least two crops at the same time on the same field, could be a promising way to enhance soil microbial diversity and functioning by increasing plant diversity in arable systems. This increase in soil diversity and functioning could then feedback on crop yield through enhanced microbial activity and nutrient mobilisation or decreased pathogen accumulation (Wang et al., 2017; Bender & van der Heijden 2015; Zhou, Yu, & Wu, 2011).

However, despite current technological advances to study soil microbes, soil microbial communities and their assembly mechanisms are still poorly understood (Van der Putten *et al*., 2013; López-Angulo *et al*., 2020). In particular, we are still unsure what soil microbes respond to - e.g. plant aboveground traits, plant root traits, plant community composition, plant functional diversity - and if these responses to the biotic environment are comparable to the effects of abiotic conditions, such as soil moisture or nutrients. Consequently, the extent to which plant–microbe interactions in soils could influence crop yield of intercropping systems is still unclear. To shed light on this subject, we conducted an extensive common garden experiment including eight annual crop species belonging to four functional groups, and forty different crop mixtures of two and four species, in which we determined soil microbial composition and measured soil respiration in addition to crop yield. Moreover, we repeated the mesocosm experiment in two different countries – i.e. in Spain and Switzerland, which differ drastically in terms of climate and soil, and with and without fertiliser application in a fully factorial design. This experimental setup allowed us to examine the following research questions: (1) How does crop diversity affect soil microbial diversity and functioning in comparison to abiotic properties? (2) Are changes in soil microbial diversity and functioning related to crop yield? (3) Are crop–soil microbe relationships and their effects on crop yield environmentally context-dependent? We hypothesized that increasing crop diversity would lead to an increase in microbial (alpha) diversity as well as changes in bacterial and fungal community compositions – e.g. increases in symbionts, decomposers, or disease suppressive microbes – and that these changes would have a positive effect on soil microbial activity (i.e. soil basal respiration) and crop yield. Finally, we hypothesized that changes in crop–microbe relationships and their impacts on crop yield would be context-dependent, with varying size effects in Switzerland and in Spain, which differ in terms of resource availability and environmental harshness.

## 2. Materials and methods

### 2.1 Study sites

The crop diversity experiment took place in outdoor experimental gardens in Zurich, Switzerland, and in Torrejón el Rubio, Cáceres, Spain. In Zurich, the garden was located at the Irchel campus of the University of Zurich (47.3961 N, 8.5510 E, 508 m a.s.l.). In Torrejón el Rubio, the garden was situated at the Aprisco de Las Corchuelas research station (39.8133 N, 6.0003 W, 350 m a.s.l.). Zurich is characterized by a temperate climate, while the Spanish site is located in a dry, Mediterranean climate. The main climatic differences during the respective growing seasons (i.e. between April and August in Switzerland and between February and June in Spain) were precipitation (587 mm in Zurich, 229 mm in Cáceres) and daily average hours of sunshine (5.7 h in Zurich, 8.8 h in Cáceres). Temperatures during the growing season did not vary substantially between the two sites: averages of daily mean, minimum and maximum temperatures were 15.8 °C, 10.9 °C and 21.1 °C in Zurich versus 15.5 °C, 9.7 °C and 21.4 °C in Cáceres. All climatic data come from the respective national meteorological archives and are average values over the years 2000 to 2018 (www.meteoschweiz.admin.ch, www.datosclima.es).

The experimental gardens were irrigated during the growing season with the aim of maintaining the above-mentioned differences in precipitation between the two sites while assuring survival of the crops during drought periods. In Spain, the automated irrigation system was configured for a dry threshold of soil moisture of 17% of field capacity, with a target of 25%. In Switzerland, the dry threshold was set at 50% of field capacity, with a target of 90% of field capacity. Whenever dry thresholds were reached (measured through PlantCare soil moisture sensors (PlantCare Ltd., Switzerland), irrigation was started and water added until reaching the target value.

Each experimental garden consisted of square plots of 0.25 m^2^ with a depth of 40 cm. Beneath 40 cm the plots were open, allowing unlimited root growth. The plots were embedded into larger beds: in Switzerland, there were 14 beds of 14×1 m, each bed containing 28 plots. In Spain, beds were 20×1 m and contained 40 plots. Inside a bed, plots were separated from each other by metal frames. Each plot was filled until 30 cm with standard, not enriched, agricultural soil coming from the local region. Therefore, soil structure and composition varied between sites and reflected the environmental histories of each site. In Spain, soil was composed of 78 % sand, 20 % silt, and 2 % clay, and contained 0.05 % nitrogen, 0.5 % carbon, 253 mg total P/kg. In Switzerland, the soil consisted of 45 % sand, 45 % silt, and 10 % clay, and contained 0.19 % nitrogen, 3.39 % carbon, and 332 mg total P/kg. Spanish and Swiss soils had a mean pH of 6.30 and 7.25, respectively.

We fertilized half of the beds with nitrogen, phosphorus and potassium at the concentration of 120 kg/ha N, 205 kg/ha P, and 120 kg/ha K. Fertilizers were applied three times, namely once just before sowing (50 kg/ha N, 85 kg/ha P, 50 kg/ha K), once when wheat was at the tillering stage (50 kg/ha N, 85 kg/ha P, 50 kg/ha K), and once when wheat was flowering (20 kg/ha N, 34 kg/ha P, 20 kg/ha K). The other half of the beds served as unfertilized controls. We randomly allocated individual beds to a fertilized or non-fertilized control treatment (Fig. S1).

### 2.2. Crop species

At each site, experimental communities were constructed with eight annual crop species. We used crop species belonging to four different phylogenetic groups with varying functional characteristics: *Triticum aestivum* (wheat, C3 grass) and *Avena sativa* (oat, C3 grass), *Lens culinaris* (lentil, legume) and *Lupinus angustifolius* (lupin, legume), *Linum usitatissimum* (flax, herb [superrosids]) and *Camelina sativa* (false flax, herb [superrosids]), and *Chenopodium quinoa* (quinoa, herb [superasterids]) and *Coriandrum sativum* (coriander, herb [superasterids]). This resulted in species combinations with different phylogenetic distances: Cereals diverged from the three other groups 160 million years ago; superasterid herbs then diverged from superrosid herbs and legumes 117 million years ago. Finally, legumes diverged from superrosid herbs 106 million years ago (timetree.org). Furthermore, we chose crop cultivars that were commercially available in Switzerland (the list of cultivars and suppliers can be found in Table S1 in SI).

### 2.3 Experimental crop communities

Experimental communities consisted of control plots with no crops, monocultures, 2- and 4-species mixtures. We planted every possible combination of 2-species mixtures with two species from different phylogenetic groups and every possible 4-species mixture with a species from each of the four different phylogenetic groups present. We replicated the experiment two times with the exact same species composition in each country. Within each country, plots were randomized within each fertilizer treatment. Each monoculture and mixture community consisted of one, two or four species planted in four rows. Two species mixtures were organized following a speciesA|speciesB|speciesA|speciesB pattern. The order of the species was chosen randomly. For 4-species mixtures, the order of the species was also randomized. Density of sowing differed among species groups and was based on current cultivation practices: 160 seeds/m^2^ for legumes, 240 seeds/m^2^ for superasterids, 400 seeds/m^2^ for cereals, and 592 seeds/m^2^ for superrosids. Seeds were sown by hand in early February 2018 in Spain and early April 2018 in Switzerland.

### 2.4 Data collection

*Soil samples:* Soil samples were collected in each plot during flowering of the crops (early May in Spain, early June in Switzerland). We took three samples per plot to a depth of 25 cm, one between each of the four plant rows of a plot, which we then pooled. These soil samples were transported cooled to the lab and sieved with a 2mm mesh size.

#### Soil activity measured as soil basal respiration

With the sieved soil samples, we then measured water content, water holding capacity, and incubated 25 g equivalent dry soil at a normalized moisture level (60% WHC) for 12 hours in airproof jars. The CO_2_ content in the headspace was measured with a LiCor LI820 once at the beginning of the incubation and once at the end. The difference between the two measures corresponded to the amount of CO_2_ that had been produced by soil microbial respiration (Curiel Yuste *et al*., 2007).

#### DNA extraction and amplification

Total nucleic acids were extracted from 250 mg of sieved soil using the DNeasy PowerSoil Kits (Qiagen, Hilden, Germany) following the supplier’s protocol. Concentration of extracted DNA was measured photospectrometrically with the QIAxpert System (Qiagen). Amplicon sequencing library construction was performed following a two-step PCR approach. The first step was performed using specific primers targeting the bacterial and archeal 16S ribosomal RNA gene (region V3-V4) and the fungal internal transcribed spacer region ITS2 using primer pairs 341F and 806R (Frey *et al*., 2016) and ITS3ngs and ITS4ngsUni (Tedersoo & Lindahl, 2016), respectively, including the sequencing primer sites of the Illumina adapters P5 (CTTTCCCTACACGACGCTCTTCCGATCT) and P7 (GGAGTTCAGACGTGTGCTCTTCCGATCT) required for the second step index PCR (Illumina Inc., San Diego, CA, USA). PCR amplification was performed in a volume of 25 µl containing 20 ng of template DNA, 1x GoTaq^®^ Flexi Buffer (Promega, Madison, WI, USA), mM MgCl_2_ (Promega), 0.4 µM of each primer (Microsynth, Balgach, Switzerland), 0.2 mM dNTPs (Promega), 0.6 mg ml^-1^ BSA (VWR, Radnor, PA, USA) and 1.25 U GoTaq^®^ G2 Hot Start Polymerase (Promega). The PCR conditions consisted of an initial denaturation at 95 °C for 5 min, 28 (16S rRNA gene) or 35 (ITS2 rrn) cycles of denaturation at 95 °C for 40 s, annealing at 58 °C for 40 s and elongation at 72 °C for 1 min, followed by a final elongation at 72 °C for 10 min. Each sample was amplified in triplicates and subsequently pooled. Pooled DNA samples were sent to the Functional Genomics Center Zurich (FGCZ, Zurich, Switzerland) for the indexing PCR. Index PCR products were purified, quantified, and pooled in equimolar ratios prior to sequencing on the Illumina MiSeq platform using the v3 chemistry (Illumina Inc.).

#### Soil moisture level

At the same time as the collection of soil samples, volumetric soil water content was measured from the soil surface to a depth of 6 cm using a ML3 ThetaProbe Soil Moisture Sensor (Delta-T, Cambridge). The measurements were taken in the center of each of the three in-between rows per plot and averaged per plot.

#### Grain yield

Grain yield of each crop species was determined in each plot when grains reached maturity (duration of crop growth from sowing to harvest given in Table 1). As time of maturity slightly varied among the different crops, we harvested species by species.

**Table 1:**
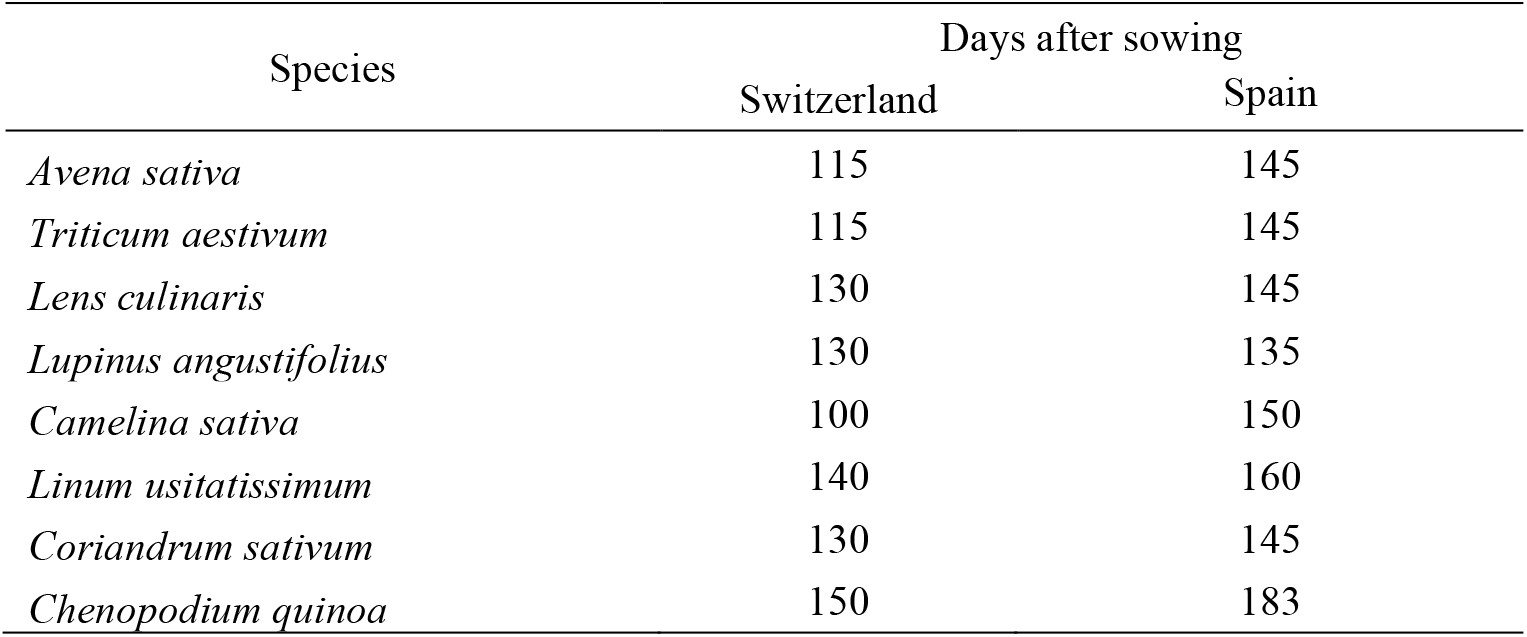
Crop growth duration in mean days after sowing from sowing until harvest for both countries and all eight species.

### 2.5 Data analyses

Due to wild birds foraging on wheat seeds in Spain, and consequently, loss of data on wheat yield, we discarded the plots that were affected by the birds foraging. A total of 341 out of 384 plots remained, 181 in Switzerland and 160 in Spain.

#### Bioinformatics

Sequences were processed using a customized pipeline largely based on VSEARCH. The main steps included paired-end read merging using the *fastq_mergepairs* algorithm implemented in VSEARCH (Edgar & Flyvbjerg, 2015); primer trimming using Cutadapt (Martin, 2011) allowing for one mismatch; removing PhiX control reads using Bowtie2 (Langmead & Salzberg, 2012); quality filtering by maximum expected error of one using the *fastq_filter* function (Edgar & Flyvbjerg, 2015) implemented in VSEARCH; dereplicating sequences using the *derep_fulllength* function in VSEARCH; delineation of sequences into amplicon sequence variants (ASVs) using the *unoise3* algorithm (Edgar, 2016) implemented as the *cluster_unoise* function in VSEARCH with an alpha of 2 and minsize of 4; removal of potentially chimeric sequences using the uchime2 algorithm (Edgar, 2016a) implemented as the *uchime3_denovo* function in VSEARCH; target verification using Metaxa2 (Bengtsson-Palme *et al*., 2015) and ITSx (Bengtsson-Palme *et al*., 2013) for the 16S rRNA gene and ITS2 sequences, respectively; mapping of the quality-filtered reads of each sample against the verified ASV sequences using the *usearch_global* function in VSEARCH; and taxonomic classification of each ASV sequence using the SINTAX classifier (Edgar, 2016b) implemented in VSEARCH with a bootstrap support of 0.8 against the SILVA database for the 16S rRNA sequences (Pruesse *et al*., 2007) and the UNITE database for the ITS2 sequences (Abarenkov *et al*., 2010). Non-fungal ITS2 sequences, as well as 16S rRNA sequences assigned to organelle structures (chloroplasts, mitochondria) were removed from the ASV table.

To remove effects of variability in sequencing read numbers, we performed 100-fold iterative subsampling of the ASV table using the function *rrarefy* from the *vegan* package in R (Oksanen *et al*., 2019; R Core Team, 2019), and subsequently computed the mean abundance for each ASV. Alpha diversity was assessed by calculating Shannon’s diversity (H) and Pielou’s evenness (J) indexes.

We used linear mixed models followed by type I analysis of variance to analyze the effects of the experimental treatment on fungal and bacterial ASV richness, H, and J, respectively. The analyses were performed for Spain and Switzerland separately. Fixed factors included fertilizing condition, crop species number (i.e. two or four crop species) nested in monoculture vs. mixtures, presence of cereal, presence of legume, presence of superrosid herb, presence of superasterid herb, as well as the interactions between them (except interactions between crop species number and monoculture vs. mixture and presence of functional group, respectively). Presences of the different functional groups were all binary factors (yes, no). Species composition and bed were set as random factors. Homogeneity of variance and normality of residues were assessed visually and with Shapiro tests (Royston, 1982).

To analyze the responses of the microbial community composition to the experimental treatments, we first performed principal coordinates analysis (PCoA) using Bray-Curtis dissimilarity on relative ASV abundance (Gower, 1966). To that end, sparse ASVs (i.e. ASVs that appeared in one or two plots only) were removed, and relative abundance was log-transformed. We first performed this analysis taking both countries together, and in a second time, per country separately. Then, we used permutational multivariate analyses of variance (PERMANOVA, (Anderson, 2001)) with Bray-Curtis dissimilarity on the previously described composition matrices (i.e. separated per country), using the function *adonis* from the *vegan* package with 999 permutations (Oksanen *et al*., 2019). Experimental factors tested included fertilization, crop diversity treatments (monocultures vs. mixtures, crop species number, presence of the different functional groups and/or species), and their interactions. Homogeneity of variance was tested using permutational analysis of multivariate dispersion (PERMDISP, (Anderson, 2006)) implemented as the *betadisper* function in *vegan* with 999 permutations. Finally, we conducted a canonical analysis of principal coordinates (CAP, (Anderson & Willis, 2003)) for each country using the function *CAPdiscrim* from the *BiodiversityR* package (Kindt & Coe, 2005).

We analysed species-specific and phylum-specific responses to the experimental factors with indicator species analyses using the *indicspecies* package (Cáceres & Legendre, 2009), calculating the point-biserial correlation coefficient, and correcting for multiple testing using the Benjamini-Hochberg method with the *p*.*adjust* function from the *stats* package (Benjamini & Hochberg, 1995).

Considering the possibilities of direct and indirect effects of the different environmental and experimental variables (fertilizer, crop species richness) on soil moisture, soil microbes, soil activity, and crop yield (measured as total grain mass per plot), we applied structural equation modelling (SEM) to our data sets, separately for Switzerland and Spain. The SEMs were built, run and evaluated with the *lavaan* package (Rosseel, 2012). We decided to fit the SEM to the Swiss and Spanish datasets separately because microbial composition in the two countries showed large differences. SEM allows to test complex a priori defined direct and indirect relationships in a unique framework and to assess the overall fit of the data to the model (Grace, 2006). For our a priori model, we considered environmental variables (fertilizer), crop species richness, soil moisture level in terms of volumetric water content, soil microbial alpha diversity measures (bacterial and fungal Shannon indexes), soil microbial beta diversity measures (coordinates from the three first axes of the principal coordinates analyses for fungi and bacteria, respectively), soil activity measured as CO_2_ flux, and total crop yield per plot. Our a priori model relating environmental and experimental factors, soil moisture, soil microbial diversity, soil activity and crop yield included the following hypotheses: (1) soil microbial diversity components (i.e. alpha diversity and community composition axes) are directly influenced by the environmental conditions (fertilizer), soil moisture, and the experimental treatment (crop species richness). (2) Soil activity can be affected directly or indirectly by crop species richness, soil moisture and fertilizer; the relationship can be indirect if effects on soil activity are mediated by effects on soil diversity measures. (3) Crop yield can also be directly or indirectly affected by crop species richness, soil moisture and fertilizer. The relationship can be indirect if effects on crop yield are mediated through changes in soil microbial diversity measures or soil activity. (4) All the soil microbial diversity measures can covary with each other.

To overcome scale differences among variables, we log-transformed crop yield before inclusion in the SEM. Path coefficients were estimated using maximum likelihood, and the model fits were tested with a chisquare goodness of fit test, a Bollen–Stine bootstrap test with 1000 bootstrap draws, a root mean square error of approximation (RMSEA) test, and the comparative fit index (CFI). A non-significant chisquare, Bollen–Stine and RMSEA test, as well as CFI values above 0.90, indicate a good fit of the model to the data (Kline, 2011).

Finally, we calculated the weighted average score of each ASV based on the previously mentioned PCoA decompositions, and subsequently investigated with linear models which bacterial and fungal genera were non-randomly distributed along the PCoA axes.

## 3. Results

After sequencing, the fungal dataset contained 9,713,470 reads delineated into 7856 ASVs. After removing all non-fungal ASVs, we obtained 6,161,288 reads assigned to 3546 ASVs. The vast majority of the removed ASVs were assigned to plants despite having sieved the soils with 2mm mesh size, which suggests low fungal biomass in our system. The bacterial dataset contained 6,394,402 reads delineated into 44136 ASVs. After removing all sequences associated with chloroplasts and mitochondria, we obtained 6,306,145 reads assigned to 43857 ASVs.

### 3.1 Soil microbial alpha diversity

All diversity metrics were consistently lower in Switzerland compared to Spain (**Error! Reference source not found**.): mean fungal ASV richness was 608 in Spain, against 324 in Switzerland (−47%), whereas fungal H was lower by 22% and fungal J by 13%. Similarly, mean bacterial ASV richness was lower by 30% (i.e., 803 versus 561 ASVs in Spain versus Switzerland); bacterial H was 16% lower and bacterial J 11% lower in Switzerland compared to Spain.

In Spain, bacterial H was marginally lower (2%) in fertilized compared to unfertilized plots (Table S2, Fig 1b). Furthermore, the presence of a superasterid herb marginally increased bacterial ASV richness (+2.5%) and decreased bacterial J (−1%).

**Figure 1.**
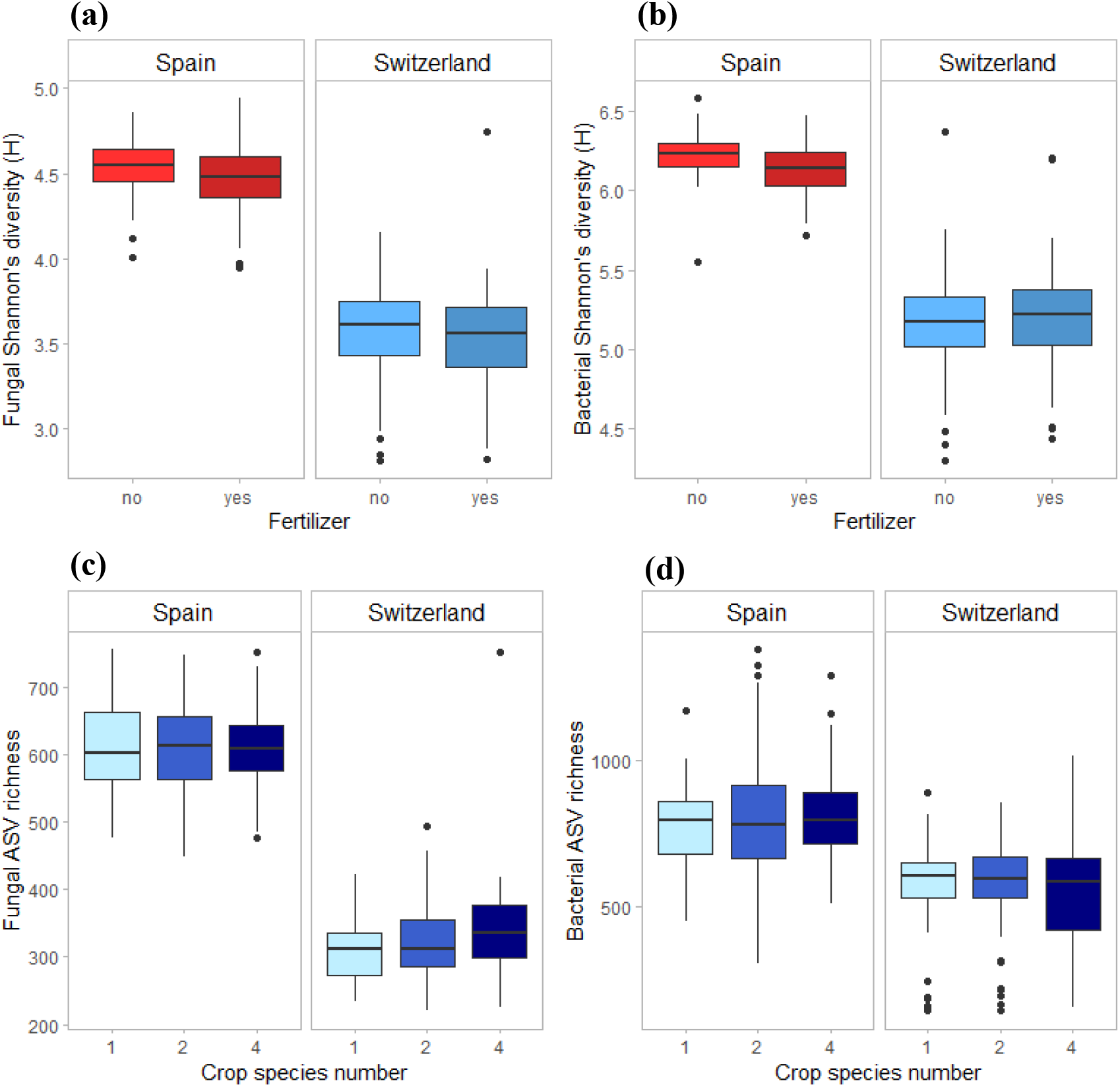
Effects of fertilizer on fungal (a) and bacterial (b) Shannon’s diversity index (H), and of crop species number on fungal (c) and bacterial (d) ASV richness in Spain and Switzerland. Horizontal lines represent the median of the data, boxes represent the lower and upper quartiles (25% and 75%), with vertical lines extending from the hinge of the box to the smallest and largest values, no further than 1.5 * the interquartile range. Data beyond the end of the whiskers are outlying and plotted individually. See Tables S2 and S3 for the results of the statistical analyses.

In Switzerland, fertilization decreased bacterial ASV richness (−16% in fertilized compared to unfertilized plots) but increased bacterial evenness (+4%) (Table S3). Fungal ASV richness marginally increased with crop species number (Fig 1c; +7% in 4-species mixtures compared to monocultures and +4% compared to 2-species mixtures), while bacterial ASV richness was higher in 2-species mixtures compared to 4-species mixtures (Fig 1d). Finally, the presence of a legume decreased fungal H (−1%) and J (−1%) in comparison to plots without legumes, while cereal presence decreased bacterial J (−2%).

### 3.2 Soil microbial community composition

Fungal and bacterial community composition differed strongly between countries (Fig. S2 of SI), with the country accounting for 66% and 52% of the variance in fungal and bacterial communities, respectively (Table S4 of SI). All other factors explained less than 1% of the remaining variance. In Spain, fungal communities were characterized by higher relative abundance of *Ascomycota, Mortierellomycota* and *Mucoromycota*, while the fungal communities in the Swiss soils had relatively more *Basidiomycota, Chytridiomycota* and *Rozellomycota* (Figs S3a and S4a of SI). Notable changes in bacterial communities between the two countries include higher relative abundance of *Planctomycetes, Actinobacteria, Acidobacteria*, and *Nitrospirae* in Spain, while in Switzerland bacterial communities showed more *Proteobacteria, Chloroflexi, Bacteroidetes*, and *Cyanobacteria* (Figs S3b and S4b of SI).

In Spain, the community composition of both fungi and bacteria was significantly affected by fertilization, presence of cereal, as well as their interaction (Table 2, Fig S5 and S6 of SI). The interaction between fertilizer and the presence of legume also had an effect on fungal communities in Spain. Fertilized plots had fungal communities with relatively more *Chytridiomycota*, while *Mortierellmycota, Mucoromycota* and *Kickxellomycota* were positively linked to unfertilized plots (Fig S7a of SI). For bacterial communities, we observed a relative increase of *Proteobacteria, Actinobacteria, Nitrospirae, Firmicutes* and *Patescibacteria* in fertilized plots, while *Planctomycetes, Acidobacteria*, and *Chloroflexi* were more abundant in unfertilized plots. *Deinococcus-Thermus* and *Fibrobacteres* increased in relative abundance in the presence of cereals, while *Nitrospirae* was associated with the absence of cereals (Fig S9a of SI).

**Table 2:** Results of the permutational analysis of variance, showing R^2^ and significance of the considered factors for bacteria and fungi in Switzerland and Spain. P-values are significant at α = 0.05; * (P < 0.05), ** (P < 0.01), *** (P < 0.001).

|  | *Fungi* |  |  |  |  |  | *Bacteria* |  |  |  |  |  |
| --- | --- | --- | --- | --- | --- | --- | --- | --- | --- | --- | --- | --- |
|  | *Spain* |  |  | *Switzerland* |  |  | *Spain* |  |  | *Switzerland* |  |  |
| | $R^2$ | $p$-value | | $R^2$ | $p$-value | | $R^2$ | $p$-value | | $R^2$ | $p$-value | |
| *Fertilizer* | **0.0471** | **0.0010** | *** | **0.0337** | **0.0010** | *** | **0.0427** | **0.0010** | *** | **0.0425** | **0.0010** | *** |
| *Monoculture vs mixture* | 0.0054 | 0.8470 |  | 0.0062 | 0.2670 |  | 0.0070 | 0.2980 |  | 0.0055 | 0.3920 |  |
| *Crop species number (2 vs 4)* | 0.0068 | 0.3820 |  | **0.0132** | **0.0020** | ** | 0.0072 | 0.2420 |  | **0.0104** | **0.0040** | ** |
| *Cereal* | **0.0156** | **0.0010** | *** | **0.0178** | **0.0010** | *** | **0.0093** | **0.0380** | * | **0.0130** | **0.0020** | ** |
| *Legume* | 0.0083 | 0.1130 |  | **0.0165** | **0.0020** | ** | 0.0079 | 0.1510 |  | **0.0124** | **0.0010** | *** |
| *Superasterid herb* | 0.0087 | 0.0640 |  | **0.0093** | **0.0320** | * | 0.0069 | 0.3050 |  | 0.0077 | 0.0660 |  |
| *Fertilizer x mono vs mixture* | 0.0065 | 0.4630 |  | 0.0051 | 0.4970 |  | 0.0059 | 0.7190 |  | 0.0077 | 0.0590 |  |
| *Fertilizer x species number* | 0.0067 | 0.4210 |  | 0.0052 | 0.4820 |  | 0.0060 | 0.6740 |  | 0.0048 | 0.6960 |  |
| *Fertilizer x cereal* | **0.0100** | **0.0290** | * | 0.0059 | 0.3150 |  | **0.0101** | **0.0240** | * | 0.0055 | 0.4010 |  |
| *Fertilizer x legume* | **0.0099** | **0.0240** | * | 0.0038 | 0.8580 |  | 0.0081 | 0.1270 |  | **0.0079** | **0.0430** | * |
| *Fertilizer x superasterid herb* | 0.0054 | 0.8190 |  | 0.0055 | 0.4070 |  | 0.0062 | 0.5980 |  | 0.0069 | 0.1200 |  |

In Switzerland, fungal and bacterial communities were affected by fertilization, crop species number, presence of cereal and presence of legume (Table 2, Fig S5 and S6 of SI). Fungal communities were also affected by the presence of superasterid herbs. The interaction of fertilization and presence of legume had an effect on bacterial communities. *Mucoromycota* were more abundant in fertilized plots, while *Ascomycota* were more present in unfertilized plots (Fig S7b of SI). For bacterial communities, we noticed an increase in *Proteobacteria, Patescibacteria, Hydrogenedentes, Bacteroidetes*, and *Cyanobacteria* in fertilized plots, while *Planctomycetes, Actinobacteria*, and *Firmicutes* were more abundant in unfertilized plots. Furthermore, there was an increase in the relative abundance of *Fibrobacteres, Verrucomicrobia*, and *Armatimonadetes* in the presence of cereals, while plots with no cereal had more *Hydrogenedentes* and *Latescibacteria* (Fig S9b of SI).

#### 3.3 Structural Equation Modelling

Our data showed overall good fit to our *a priori* SEM in Spain and Switzerland: chisquare = 3.828 (Spain) and 4.939 (Switzerland); P(chisquare) = 0.281 (Spain) and 0.173 (Switzerland); P(Bollen–Stine Bootstrap) = 0.459 (Spain) and 0.464 (Switzerland); RMSEA = 0.046 (Spain) and 0.064 (Switzerland); P(RMSEA) = 0.415 (Spain) and 0.321 (Switzerland); CFI = 0.999 (Spain) and 0.997 (Switzerland).

In Spain, soil diversity measures were mostly influenced by fertilizer and soil moisture (Fig. 2a). Soil microbial activity was directly influenced by soil moisture, fungal PCoA 2 and bacterial PCoA 2. Crop yield was only linked to bacterial PCoA axis 2. Bacterial PCoA axis 2 was associated with a response of *Bacilli, Actinobacteria* and *Acidobacteria*, among others (Fig S11 of SI), which therefore correlated with an increase in crop yield in our study. Fertilization had an indirect positive effect on crop yield via changes in bacterial PCoA axis 2.

**Figure 2.**
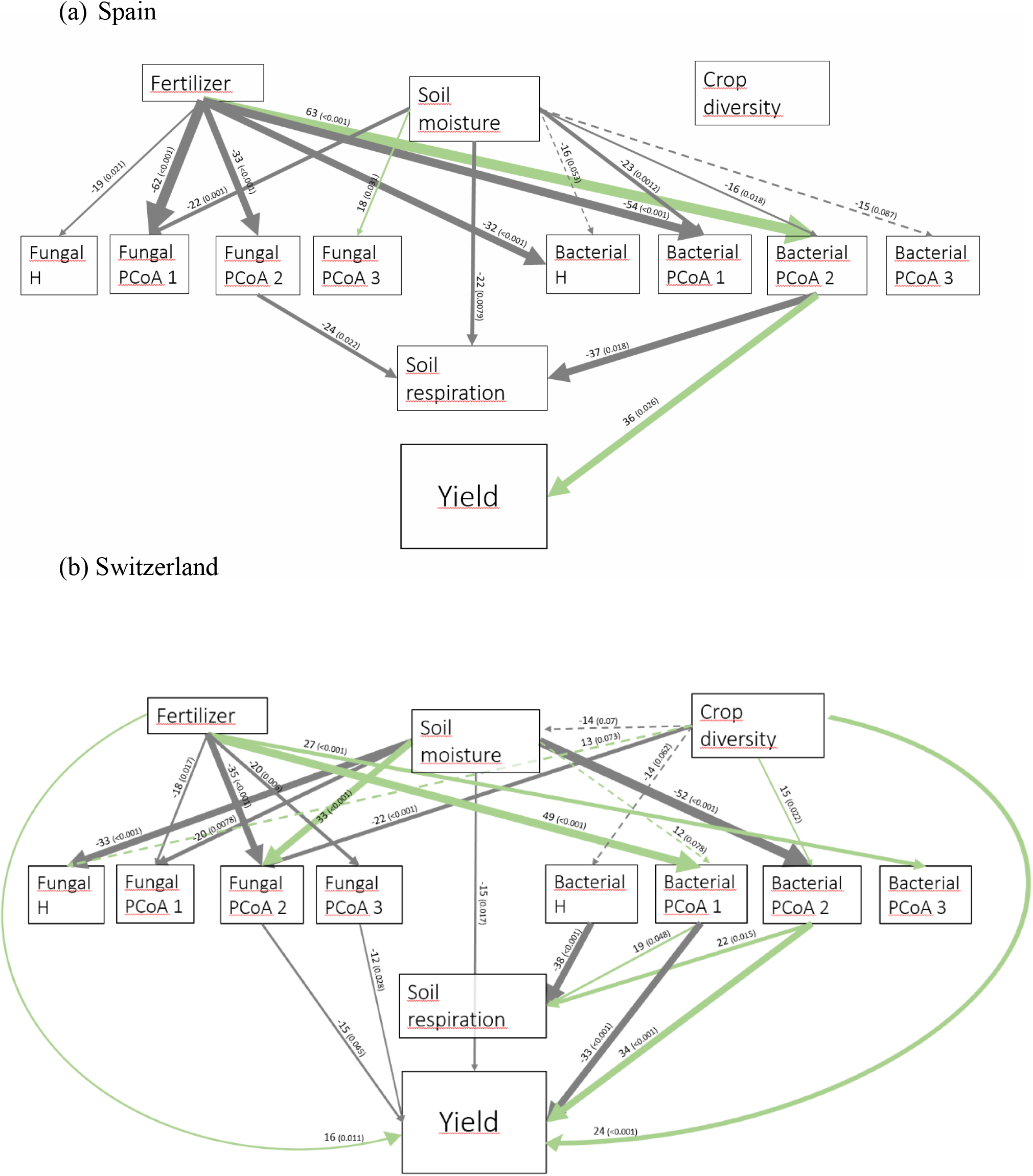
Structural Equation Modelling showing the relationships between crop diversity, fertilizer, soil moisture, soil activity, and bacterial and fungal diversity measures in Spain (a) and Switzerland (b). H: Shannon’s diversity. Only significant (solid line) and marginally significant (dashed line) relationships are shown. Width of arrows are proportional to the strength of the standardized path coefficients indicated by the numbers above the arrows. The numbers in brackets indicate associated p-values. Colours of the arrows show positive (green) and negative (grey) effects. Residuals correlations are not shown.

In Switzerland, fertilizer and soil moisture also significantly influenced almost all the soil diversity measures (Fig. 2b). Moreover, crop diversity had a direct effect on soil moisture, bacterial and fungal H, fungal PCoA axis 2, and bacterial PCoA axis 2. Soil microbial activity was negatively affected by bacterial H, and positively by the first and second axes of bacterial PCoA. Crop yield was directly affected by some soil diversity measures (fungal PCoA axes 1 and 2, bacterial PCoA axes 1 and 2), and also by crop diversity, fertilizer and soil moisture. Two-species and four-species mixtures showed an increase in average yield of 43% and 102% compared to monocultures, respectively, while fertilization increased yield by 13%.

Crop diversity indirectly affected crop yield in Switzerland through five different pathways. Firstly, there were two indirect positive effects through the second axis of fungal PCoA; fungi genera associated with negative coordinates along the PCoA axis 2 included for instance *Kondoa, Serendipita, Mucor*, or *Leucospiridium*, among others (Fig S10 of SI). The presence of these fungal genera is therefore linked to an increase in crop yield. Secondly, there were two positive effects of crop diversity on crop yield mediated by the second axis of bacterial PCoA. Bacterial PCoA axis 2 is related to a response of *Nitriliruptoria, Chloroflexia, Acidimicrobia*, and *Actinobacteria*, among others (Fig S12b of SI); these bacteria thus form another potential group of yield promoting soil microbes in our study. Finally, crop diversity indirectly increased crop yield through the first axis of bacterial PCoA. Bacterial genera associated with negative coordinates along the PCoA axis 1 include *Nitriliruptoria, Actinobacteria, Acidimicrobia, Chloroflexia*, and *Bacilli*, among others (Fig S12a of SI), i.e. a third group of yield promoting soil microbes in our study. Fertilization had an indirect positive effect on crop yield, via changes in fungal PCoA axes 2 and 3, but also an indirect negative effect on crop yield, mediated by changes in bacterial PCoA axis 1.

## 4. Discussion

Our study shows that crop diversity and composition, as well as fertilisation, can affect soil microbial community composition, and that these compositional changes go along with changes in crop yield. In particular, the data suggests that in Switzerland, the positive effects of increasing crop diversity on crop yield are partially mediated by changes in microbial composition, notably through a response of plant-growth promoting bacteria. However, as expected, changes in soil microbes do not fully explain yield variations and thus other must play an important role in increasing crop yield in mixtures. Furthermore, the effects of crop diversity on microbial communities remained small compared to the effects of fertilisation. In Spain, where soil resources were limited and both crop and soil communities experienced more stressful environmental conditions than in Switzerland, we did not observe any significant responses of soil microbes to crop diversity. This context-dependence of crop diversity effects on soil microbes and ecosystem functioning may be explained by differences in abiotic factors, such as moisture, or nutrient availability.

### Crop species number effects on microbes

Our first hypothesis was that an increase in crop diversity would lead to an increase in microbial alpha diversity as well as changes in soil microbial community composition. We did observe an increase in fungal ASV richness in response to crop diversity, but this effect was only marginally present in Switzerland. This is consistent with results from studies in natural environments, where plant species richness positively correlated with fungal richness (Yang *et al*., 2017). There was no effect of crop diversity on any of the bacterial alpha diversity measures; however, in both countries, we observed changes in bacterial and fungal beta diversity in response to crop diversity and crop composition (Table 2). In Switzerland notably, we observed a shift in soil fungal and bacterial composition between 2-species mixtures and 4-specdireies mixtures. This is in agreement with previous research demonstrating that plant diversity did not correlate with microbial alpha diversity, but it did with beta diversity (Grüter *et al*., 2006; Prober *et al*., 2015). Moreover, the presence of a cereal among the crop species induced changes in fungal and bacterial soil communities in both countries, with a notable increase in abundance of *Fibrobacteres, Armatimonadetes*, and *Verrucomicrobia*. The two latter groups have commonly been found in the rhizosphere of wheat or oat, even though little is known about their functions (Rascovan *et al*., 2016; Dai *et al*., 2020; Lupwayi *et al*., 2020). The response of *Fibrobacteres* - which are mostly involved in the degradation of plant-based cellulose (Gupta, 2004) - to the presence of a cereal has been scarcely mentioned in previous research (Mahoney *et al*., 2017).

### Soil microbes partially explain the positive effects of intercropping on yield

Results from our SEM in Switzerland suggest that the changes in soil microbial communities induced by crop diversity further affected ecosystem functioning and services, such as soil microbial activity and crop yield. We had previously shown that there was a positive effect of crop species richness on crop yield in Switzerland, demonstrating a positive diversity– productivity relationship (Chen *et al*., 2020; Engbersen *et al*., 2020). Here, we provide evidence that these positive effects of intercropping on crop yield can partially be explained by changes in microbial communities. Indeed, increasing crop diversity was linked to an increase in yield through five indirect pathways mediated by changes in fungal or bacterial composition (Fig 2b). We noticed that those changes in microbial composition associated with increases in crop yield were characterized by a response of potentially beneficial bacteria such as plant growth-promoting rhizobacteria (PGPRs). For instance, this was the case of *Actinobacteria* (Fig S11, S12 of SI), which are known to be symbionts of plants as well as saprophytes (Bhattacharyya & Jha, 2012). They also play a role as biocontrol agents against a range of pathogenic fungi and promote plant growth (Merzaeva & Shirokikh, 2006; Franco-Correa *et al*., 2010). Some examples include *Glycomyces, Agromyces, or Nocardioides*, which are genera of PGPRs (Qin *et al*., 2011; Hamedi & Mohammadipanah, 2014), and *Streptomyces*, known to associate with wheat and possessing anti-fungal properties (Conn & Franco, 2004). Other potential PGPRs that were found to positively correlate with crop yield in our study include *Pseudomonas, Burkholderia, Bacillus*, and *Massilia* (Fig S11, S12 of SI). *Pseudomonas* has been shown to promote plant growth as well as inhibit pathogenic fungi (Gray & Smith, 2005; Melo *et al*., 2016). *Burkholderia* and *Bacillus* can be involved in phosphate solubilisation (Dimkpa *et al*., 2009; Zhao *et al*., 2014) and symbiotic associations with wheat (Shaharoona *et al*., 2007; Moreno-Lora *et al*., 2019). Finally, *Massilia* has been shown to correlate with an increase in plant biomass for alfalfa and soybean (Xiao *et al*., 2017). Some bacteria involved in nitrogen cycling also positively correlated with the PCoA coordinates leading to an increase in crop yield in Switzerland, such as *Nitrosospira –* ammonia-oxidizing genus (Fierer *et al*., 2007), and *Nitrolancea* – nitrite-oxidizing genus (Daims *et al*., 2016).

When looking at fungi, our results showed that in Switzerland crop yield was positively associated with fungal genera including *Claroideoglomus, Myrmecridium, Serendipita, Mucor*, as well as several yeasts (e.g. *Torula, Kondoa, Leucosporidium)* (Fig S10). *Claroideoglomus* belongs to the *Glomerales* order, which are biotrophic mutualists and can establish arbuscular mycorrhizal networks with wheat (Dai *et al*., 2014). *Myrmecridium* and *Mucor* are saprophytes, involved in decomposition of organic matter and nutrient cycling (Schlatter *et al*., 2018), while *Serendipita* are plant growth promoting fungi which have been shown to have beneficial effects on many plants, including wheat (Singhal *et al*., 2017). Finally, yeasts have also been suggested as potential bio-agents and plant growth promoters (Mukherjee *et al*., 2020); furthermore, they are a nutrient source for some bacteria and contribute to essential soil ecological processes such as mineralization of organic matter (Botha, 2011). Among the fungal genera associated with positive PCoA coordinates, i.e. lower crop yield, we found a few plant pathogens, such as *Protomyces*, causing stem gall disease in coriander (Khan & Parveen, 2018), and *Pyrenochaeta*, a parasite for plant roots (Aragona *et al*., 2014). Our study thus suggests that in Switzerland, increasing crop diversity leads to changes in microbial communities that enhance the presence of beneficial microbes and reduce pathogen loads, which may promote plant growth and contribute to the observed increase in crop yield.

### The importance of microbial communities is relative

While changes in soil microbial composition explained part of the positive effects of crop diversity on crop yield, there were also direct effects of intercropping on yield (i.e. not mediated by microbes) in Switzerland. This demonstrates that other mechanisms must play a role in increasing crop productivity in diverse mixtures. These processes could include a complementary use of resources, such as nutrient or light partitioning (Jesch *et al*., 2018; Engbersen *et al*., 2020), trait differentiation (Cadotte, 2017), or changes in crop–weed interactions (Stefan *et al*., 2020).

Moreover, our study highlights the context-dependency of crop diversity effects on soil microbes: indeed, abiotic factors such as fertilization, climate, soil moisture or soil texture were more important drivers of soil microbial communities than crop diversity. This is consistent with previous research indicating that soil microbial community composition mainly depends on soil moisture, temperature, and organic matter contents, and that these factors are often more important than crop diversification (Ren *et al*., 2018; Guo *et al*., 2020). In our case, in Switzerland, the effects of soil moisture and fertilization on microbial composition were indeed up to three times stronger than the effect of crop diversity (Fig 2b). Interestingly, fertilization in Switzerland seemed to have conflicting indirect effects on yield: on the one hand, it promoted fungal communities that were beneficial for crop yield, such as *Mucoromycota* (Fig S7b), but on the other hand, it also favoured bacteria that were not positively associated with crop yield, such as *Proteobacteria*. Bacterial phyla responding to mineral fertilization included *Proteobacteria, Hydrogenedentes, Bacteroidetes*, and *Cyanobacteria* (Fig S8b). This is consistent with previous studies demonstrating that the relative abundance of *Proteobacteria* and *Bacteroidetes* generally increase under high-N conditions (Fierer *et al*., 2012). *Bacteroidetes* and many proteobacterial groups have been identified as copiotrophic taxa, which tend to thrive in resource-rich environments (Fierer *et al*., 2007; Ramirez *et al*., 2010). On the contrary, *Actinobacteria*, which include many PGPRs, was significantly more abundant in unfertilized plots (Fig S8b). Therefore, we might hypothesize that high-N conditions would favour the growth of *Proteobacteria* at the expense of *Actinobacteria* (Lupwayi *et al*., 2018), thereby limiting the abundance of plant beneficial bacteria.

In Spain, soil microbes responded to soil moisture and fertilisation but did not respond to crop diversity. We propose several reasons for this lack of crop–microbe relationship in Spain. First, soil texture in Spain was dominated by sand (78%) with only a small proportion of clay (2%), unlike Switzerland where sand and silt were in equal proportions (45%) with a higher clay content (10%). Higher clay and silt content usually protect and favour microbial biomass (Bach *et al*., 2010) through larger aggregates (Wick *et al*., 2009), greater water holding capacity (Voroney, 2007), and increased nutrient retention (Knops & Tilman, 2000). Secondly, the climate in Spain is more arid than in Switzerland, which led to lower soil moisture and increased water stress for plant and soil communities. Finally, soil nutrient content was lower in Spain. Spanish soil communities thus experienced greater abiotic stress than in Switzerland, which may be why they primarily responded to soil moisture or fertilization – factors that can directly alleviate their stress – rather than crop diversity. Furthermore, there was almost no link between microbial communities and yield in Spain; in particular, fungal communities did not have any effect on crop yield, while only one axis of the bacterial composition positively affected yield

in response to fertilization (Fig 2a). This decoupling between yield and soil microbes suggests that crop productivity in Spain might be limited by other, more impactful factors that would overshadow any potential effects of microbes, such as nutrient or water availability. This is consistent with a recent study by Adomako (2020), in which they found that the positive effects of soil microbes on plant biomass were stronger under high nutrient and water availability compared to low resource conditions. Indeed, in conditions of nutrient stress, microbial communities may compete for the limited available nutrients (Ho & Chambers, 2019). If the amount of nutrients is insufficient, it is possible that microbes cannot exchange nutrients for carbon with the plants anymore, which can lead to negative plant–soil feedbacks (Petermann *et al*., 2008; in ‘t Zandt *et al*., 2019). Furthermore, nutrient or water limitation, by restricting plant growth, might also reduce the amount of carbon provided by the plants to the microbial communities (Harrison & Bardgett, 2010). This reduction in plant–soil feedbacks raises important questions on the influence of soil microbes for plant growth in resource-limited environments, and calls for further investigation to identify the limits of the role of soil microbes for increasing productivity in agricultural systems.

In conclusion, our study shows that under favourable abiotic conditions, crop diversity does influence soil microbial community composition and that these changes partially explain the positive effects of intercropping on yield, notably through an increase in the presence of potential plant growth-promoting bacteria. However, these effects are relative and less important than changes in abiotic factors, such as the addition of mineral fertilizer. Furthermore, these processes are context-dependent: in stressful environments where resources are limited, soil microbial communities do not respond to crop diversity, and their effect on crop yield is weaker. This research thus suggests that soil microbial communities can play a beneficial role in intercropped systems, but also reflects on the relative importance of microbial communities compared to abiotic factors for increasing crop productivity, thereby highlighting the context-dependency of crop–microbe relationships.

## 5. Acknowledgments

L.S. and C.S. conceived the study with input from M.H., J.S. and N.E. L.S., N.E. and C.S. collected the data; L.S. assembled and analyzed the data with the help of C.S and M.H.; L.S., M.H., and C.S. wrote the paper. All authors discussed data analyses and results.

We are grateful to Elisa Pizarro Carbonell, Carlos Barriga Cabanillas, Anja Schmutz, Roman Hüppi and Simon Baumgartner for their help with the field experiment, Manon Longepierre for her mentoring and help with the DNA lab work and analyses, Matti Barthel for his help with the soil respiration method, and Barryette Oberholzer for her help with the DNA lab work. We thank Maria Domenica Moccia at the Functional Genomics Center Zurich

(FGCZ) for optimizing the sequencing protocol and providing the sequencing service on the Illumina MiSeq platform. We also thank the Aprisco de Las Corchuelas Field Station and the University of Zurich for the use of their fields. The study was funded by the Swiss National Science Foundation (PP00P3_170645).

## Notes

### Competing Interest Statement

The authors have declared no competing interest.

