## Supplementary Information for "Positive effects of crop diversity on productivity driven by changes in soil microbial composition"

### Supporting Information

**Table S 1.** List of crop species cultivars and their suppliers in Switzerland.

| Species | Cultivar | Supplier |
|---|---|---|
| *Avena sativa* | Canyon | Sativa Rheinau |
| *Triticum aestivum* | Fiorina | DSP, Delley |
| *Coriandrum sativum* | Indian | Zollinger Samen, Les Evouettes |
| *Chenopodium quinoa* | n.a. | Artha Samen, Münsingen |
| *Lupinus angustifolius* | Boregine | Aspenhof, Wilchingen |
| *Lens culinaris* | Anicia | Agroscope, Reckenholz |
| *Camelina sativa* | n.a. | Zollinger Samen, Les Evouettes |
| *Linum usitatissimum* | Lirina | Sativa Rheinau |

**Figure S1**. Schematic representation of the mesocosm experiments. The fertilizer and crop diversity treatments were fully randomized within the two countries.

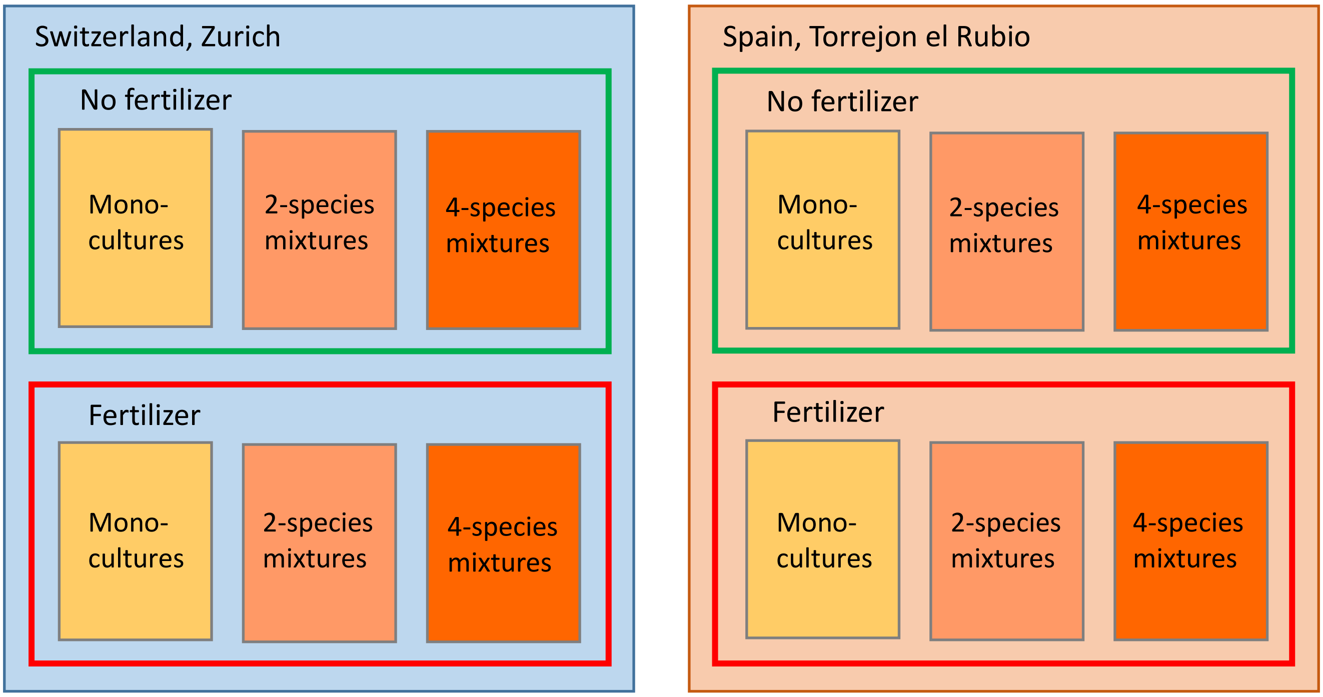

**Table S2**. Results of mixed effects ANOVA testing environmental factors, crop diversity and functional group presence on fungal and bacterial Shannon's index (H), ASV richness, and Pielou's evenness index (J) in Spain. Numbers between brackets indicate the p-value.
P-values are significant at α = 0.05 and highlighted in bold.

| | *F-value H fungi* | *F-value ASV richness fungi* | *F-value J fungi* | *F-value H bacteria* | *F-value ASV richness bacteria* | *F-value J bacteria* |
|---|---|---|---|---|---|---|
| *Fertilizer* | 3.0221 (0.11) | 3.0887 (0.11) | 1.0822 (0.35) | 4.4977 (0.057) | 2.5942 (0.13) | 1.3684 (0.26) |
| *Mono_mix* | 0.0258 (0.87) | 0.222 (0.64) | 0.1003 (0.75) | 0.0219 (0.88) | 0.3531 (0.55) | 0.0433 (0.84) |
| *Crop species number* | 0.3273 (0.57) | 0.0368 (0.85) | 0.422 (0.52) | 1.0894 (0.30) | 0.0076 (0.93) | 0.0987 (0.75) |
| *Cereal* | 0.2246 (0.64) | 0.2071 (0.65) | 0.4464 (0.51) | 0.8391 (0.36) | 1.5679 (0.21) | 0.6699 (0.41) |
| *Legume* | 2.1066 (0.15) | 0.6455 (0.42) | 1.7165 (0.19) | 0.927 (0.34) | 0.4778 (0.49) | 0.0294 (0.86) |
| *Superasterid herb* | 0.0829 (0.77) | 0.3017 (0.58) | 0.0066 (0.94) | 2.5946 (0.11) | 3.7348 (0.056) | **5.892 (0.017)** |
| *Fertilizer x mono_mix* | 0.0062 (0.94) | 1.2457 (0.27) | 0.182 (0.67) | 0.0012 (0.97) | 0.0765 (0.78) | 1.0108 (0.32) |
| *Fertilizer x crop sp.number* | 0.0634 (0.80) | 0.2356 (0.63) | 0.2357 (0.63) | 0.2191 (0.64) | 0.0132 (0.91) | 1.0014 (0.32) |

**Table S3**. Results of mixed effects ANOVA testing environmental factors, crop diversity and functional group presence on fungal and bacterial Shannon's index (H), ASV richness, and Pielou's evenness index (J) in Switzerland. Numbers between brackets indicate the p-value.
P-values are significant at α = 0.05 and highlighted in bold.

| | *F-value H fungi* | *F-value ASV richness fungi* | *F-value J fungi* | *F-value H bacteria* | *F-value ASV richness bacteria* | *F-value J bacteria* |
|---|---|---|---|---|---|---|
| *Fertilizer* | 2.3161 (0.13) | 2.8237 (0.12) | 0.0541 (0.82) | 0.0027 (0.96) | **7.0037 (0.022)** | **24.8374 (0.00060)** |
| *Mono_mix* | 1.4202 (0.24) | 2.0425 (0.15) | 1.2216 (0.27) | 1.8163 (0.19) | 0.0377 (0.85) | 2.3764 (0.13) |
| *Crop species number* | 2.1966 (0.15) | 2.964 (0.087) | 1.9326 (0.17) | 2.8809 (0.097) | **4.3801 (0.038)** | 0.0098 (0.92) |
| *Cereal* | 0.3229 (0.57) | 0.7977 (0.37) | 1.9207 (0.17) | 2.5043 (0.12) | 0.2807 (0.60) | **5.547 (0.020)** |
| *Legume* | **4.4919 (0.04)** | 3.7824 (0.054) | **5.5246 (0.02)** | 2.1157 (0.15) | 1.2589 (0.26) | 2.1281 (0.15) |
| *Superasterid herb* | 0.4028 (0.53) | 0.3254 (0.57) | 1.8028 (0.18) | 0.0072 (0.93) | 0.575 (0.45) | 1.9106 (0.17) |
| *Fertilizer x mono_mix* | 0.5653 (0.45) | 1.0359 (0.31) | 0.7852 (0.38) | **4.3963 (0.038)** | 0.5868 (0.44) | 0.4237 (0.52) |
| *Fertilizer x crop sp.number* | 0.1705 (0.68) | 0.9682 (0.33) | 0.1114 (0.74) | 1.1454 (0.29) | 0.9522 (0.33) | 0.1176 (0.73) |

**Figure S1**. Principal Coordinates Analysis plot of fungal (left) and bacterial (right) communities. Spanish samples are represented in red, while Swiss samples are represented in blue.

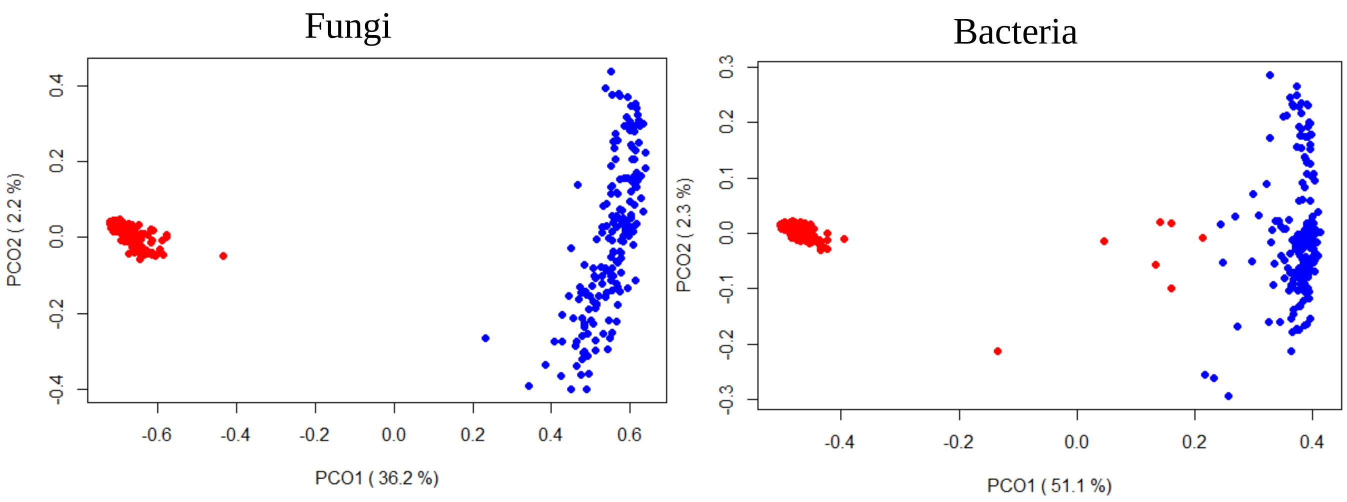


**Table S4**. Results of the permutational analyses of variance for both countries together, showing F-value, $R^2$, p-value, and significance of the considered factors.

P-values are significant at $\alpha = 0.1$; · ($P < 0.1$), * ($P < 0.05$), ** ($P < 0.01$), *** ($P < 0.001$).

| | *F_fungi* | *R2_fungi* | *Pr(>F)_fungi* | *Fungi* | *F. bact* | *R2_bact* | *Pr(>F)_bact* | *Bact* |
|---|---|---|---|---|---|---|---|---|
| *Country* | 646.81 | 0.65873 | 0.001 | *** | 357.32 | 0.51927 | 0.001 | *** |
| *Fertilizer* | 7.19 | 0.00732 | 0.002 | ** | 6.83 | 0.00993 | 0.002 | ** |
| *Mono vs mixtures* | 1.13 | 0.00115 | 0.273 | | 1.1 | 0.00159 | 0.261 | |
| *Crop species number (2 / 4)* | 1.87 | 0.00191 | 0.142 | | 1.53 | 0.00222 | 0.17 | |
| *Cereal* | 2.97 | 0.00302 | 0.057 | . | 1.79 | 0.0026 | 0.111 | |
| *Legume* | 2.44 | 0.00249 | 0.08 | . | 1.74 | 0.00252 | 0.115 | |
| *Superasterid herb* | 1.86 | 0.00189 | 0.118 | | 1.12 | 0.00162 | 0.283 | |
| *Country x fertilizer* | 5.69 | 0.00579 | 0.003 | ** | 7.08 | 0.01028 | 0.001 | *** |
| *Country x (mono vs mix)* | 0.86 | 0.00087 | 0.398 | | 0.98 | 0.00143 | 0.325 | |
| *Country x crop species number* | 1.71 | 0.00175 | 0.153 | | 1.34 | 0.00195 | 0.206 | |
| *Fertilizer x (mono vs mix)* | 0.91 | 0.00092 | 0.364 | | 1.14 | 0.00166 | 0.259 | |
| *Fertilizer x crop species number* | 1.1 | 0.00112 | 0.279 | | 0.87 | 0.00127 | 0.378 | |
| *Country x cereal* | 2.75 | 0.0028 | 0.054 | . | 1.76 | 0.00256 | 0.121 | |
| *Country x legume* | 2.12 | 0.00215 | 0.089 | . | 1.55 | 0.00226 | 0.138 | |
| *Country x superasterid herb* | 1.23 | 0.00125 | 0.25 | | 1.27 | 0.00185 | 0.217 | |
| *Fertilizer x cereal* | 1.3 | 0.00132 | 0.238 | | 1.28 | 0.00186 | 0.222 | |
| *Fertilizer x legume* | 1.02 | 0.00104 | 0.301 | | 1.33 | 0.00193 | 0.187 | |
| *Fertilizer x superasterid herb* | 1.12 | 0.00114 | 0.265 | | 1.09 | 0.00158 | 0.276 | |
| *Country x fertilizer x (mono vs mix)* | 0.96 | 0.00097 | 0.319 | | 1.08 | 0.00157 | 0.289 | |
| *Country x fertilizer x crop species number* | 0.88 | 0.00089 | 0.388 | | 0.93 | 0.00135 | 0.355 | |

**Figure S3:** Taxonomical abundance graph showing the relative proportion of the most abundant phyla of fungi (a) and bacteria (b).

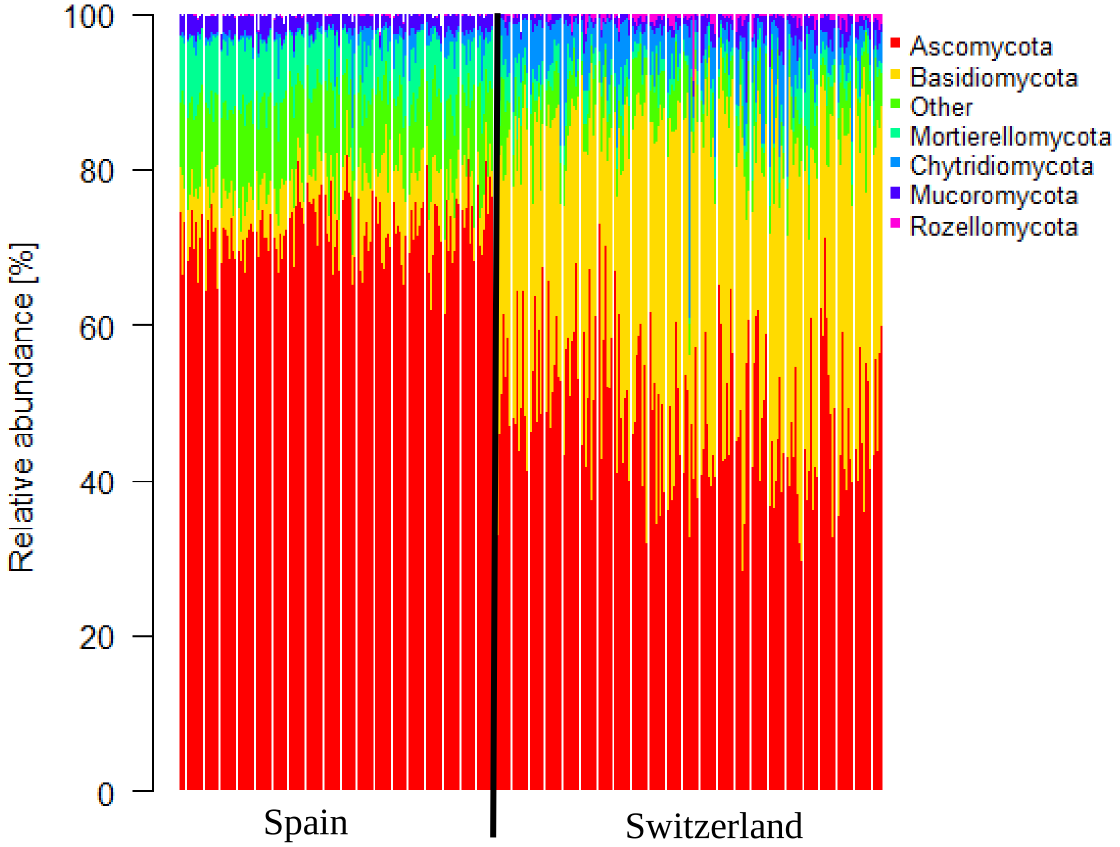


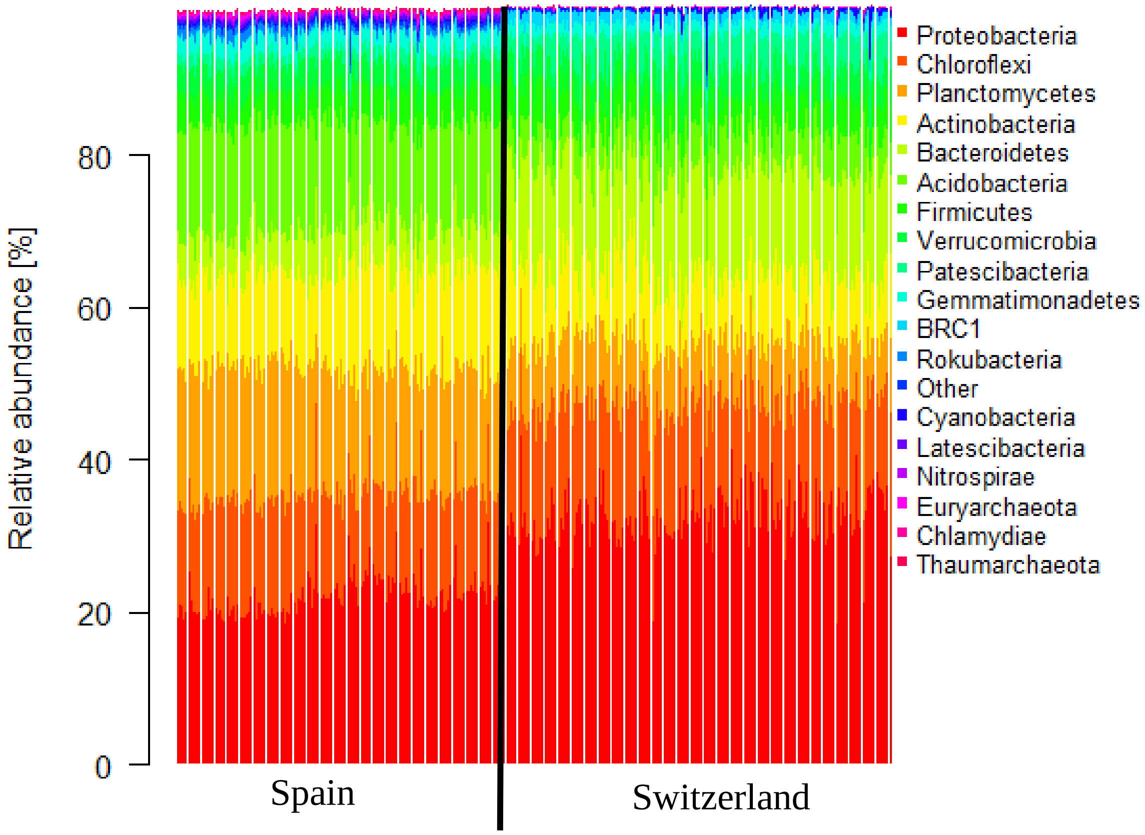

**Figure S4:** Differential abundance graph of fungal (a) and bacterial (b) phyla in the two different countries (Spain vs Switzerland).

A: percent abundance; r: point-biserial correlation coefficient; P.bh: Benjamini-Hochberg-corrected p-value.

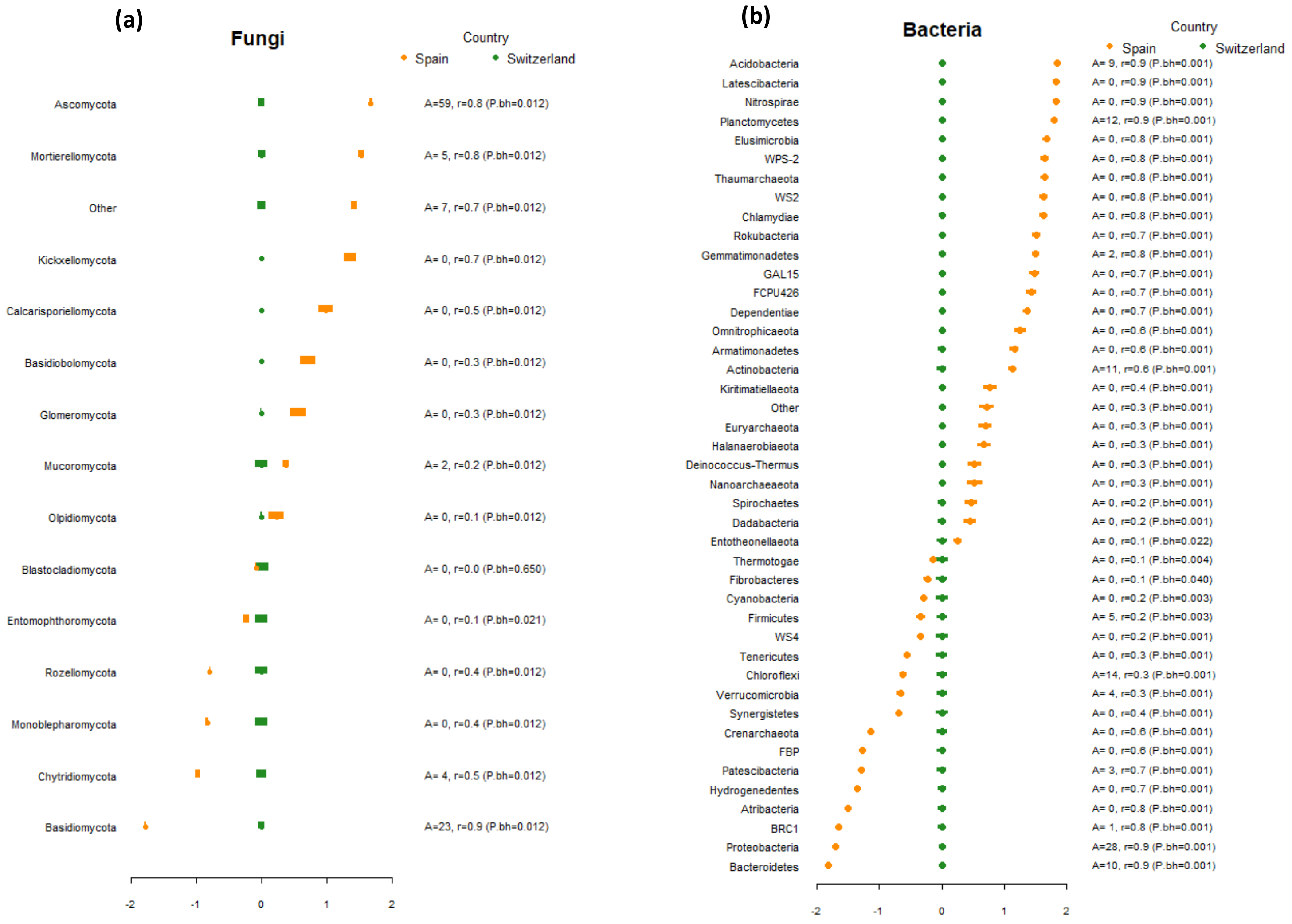

**Figure S5.** Constrained ordination plots showing changes in fungal communities in response to fertilizer (first line), cereal (second line), and crop species number (third line) in Spain (left panel) and in Switzerland (right panel). The circles represent the standard deviation of point scores of the ordination.

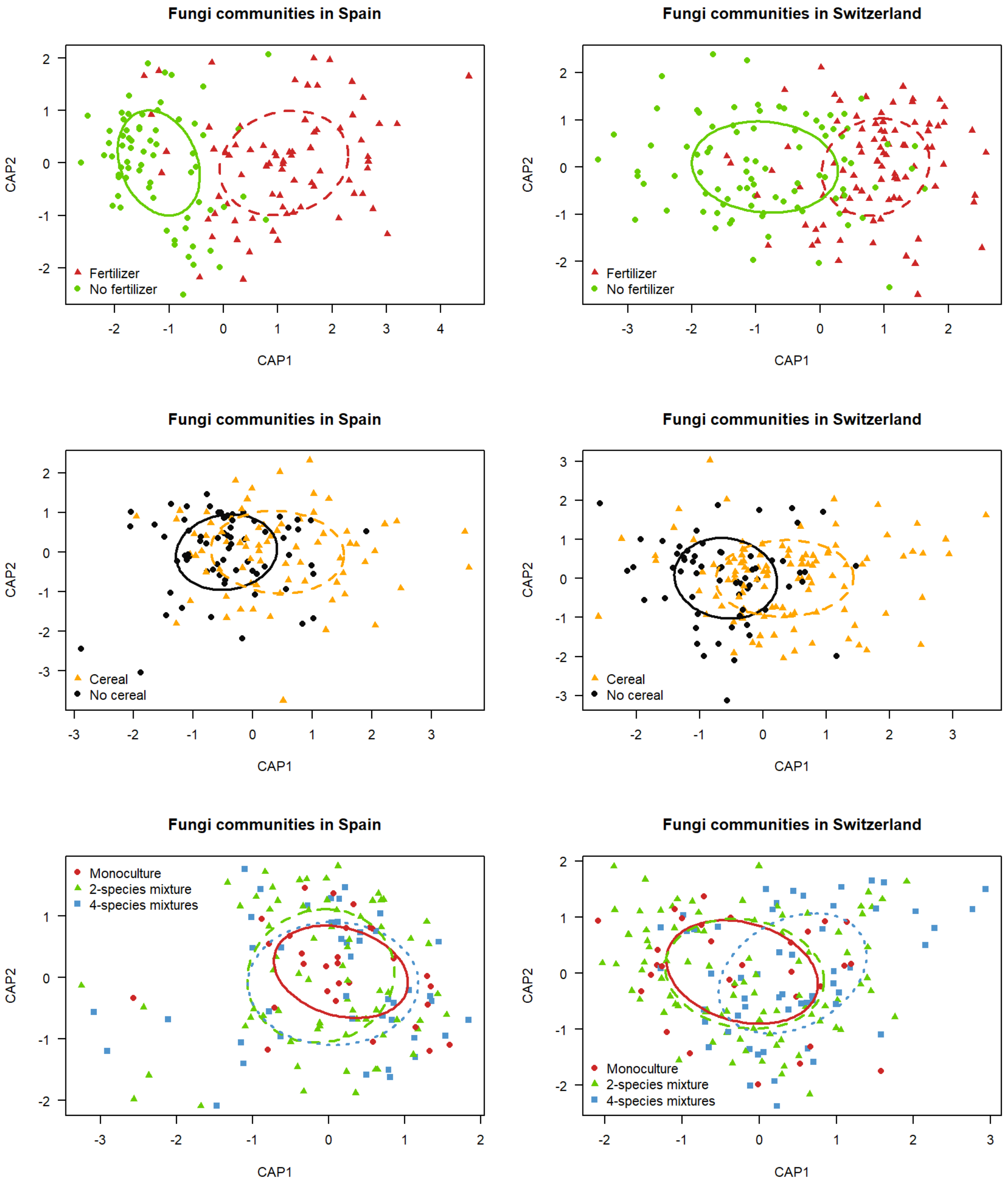

**Figure S6:** Constrained ordination plots showing changes in bacterial communities in response to fertilizer (first line), cereal (second line), and crop species number (third line) in Spain (left panel) and in Switzerland (right panel). The circles represent the standard deviation of point scores of the ordination.

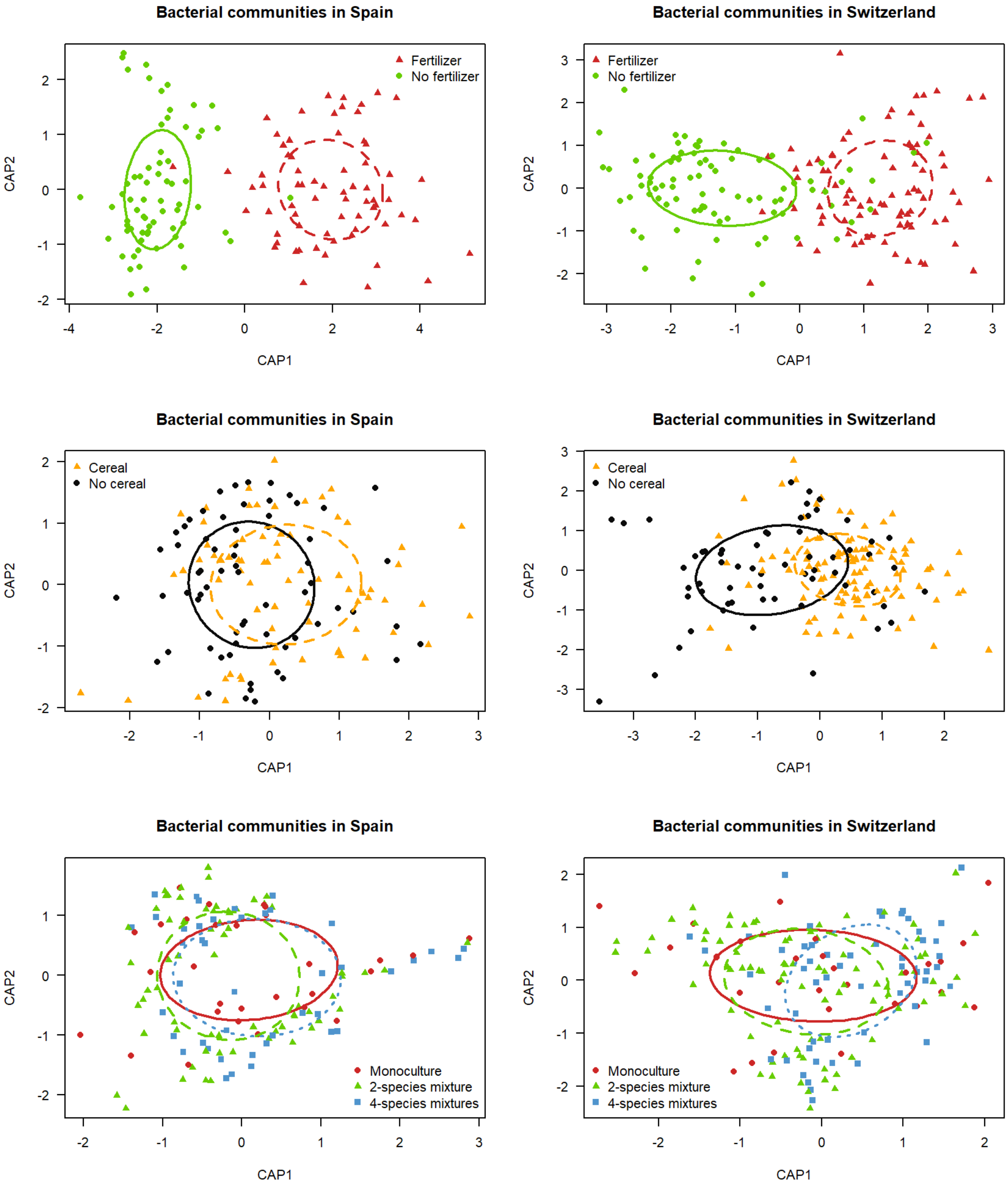

**Figure S7:** Differential abundance graph of fungi phyla under different fertilizing conditions in Spain (a) and Switzerland (b), and with or without cereal in Spain (c) and Switzerland (d).

A: percent abundance; r: point-biserial correlation coefficient; P.bh: Benjamini-Hochberg-corrected p-value.

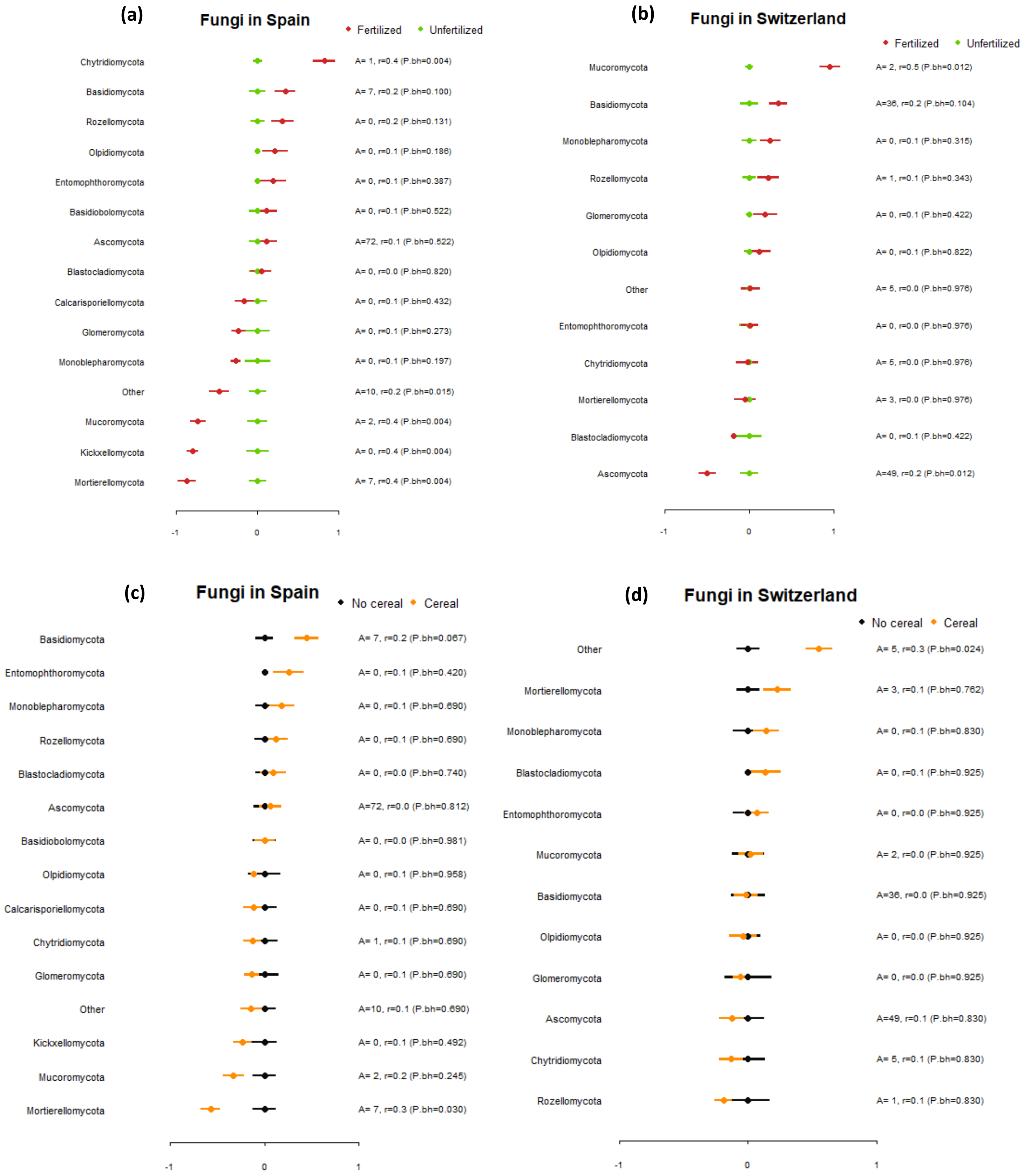

**Figure S8:** Differential abundance graph of bacterial phyla under different fertilizing conditions in Spain (a) and Switzerland (b).

A: percent abundance; r: point-biserial correlation coefficient; P.bh: Benjamini-Hochberg-corrected p-value.

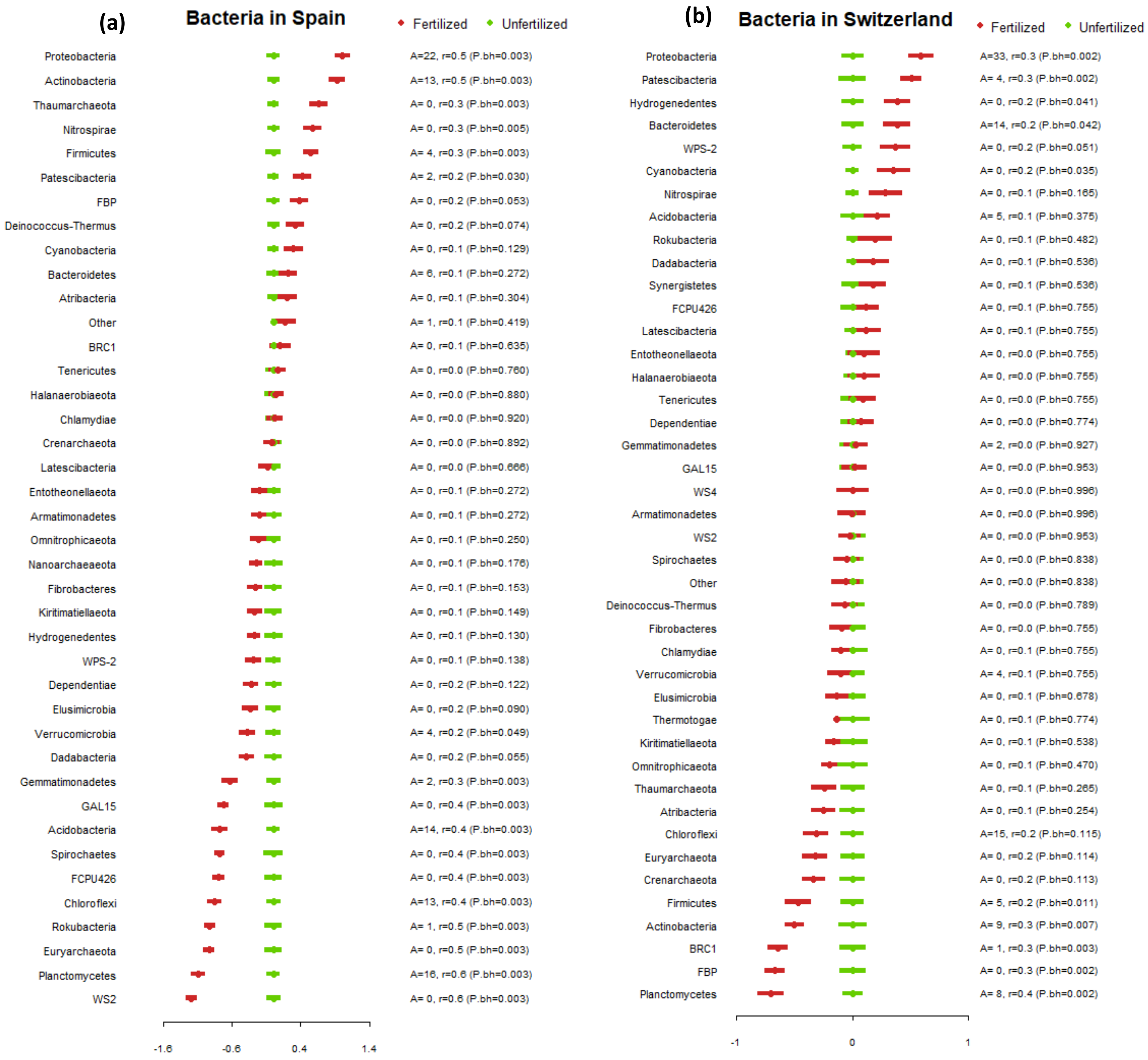

**Figure S9:** Differential abundance graph of bacterial phyla in Switzerland with or without cereal in Spain (a) and Switzerland (b).

A: percent abundance; r: point-biserial correlation coefficient; P.bh: Benjamini-Hochberg-corrected p-value.

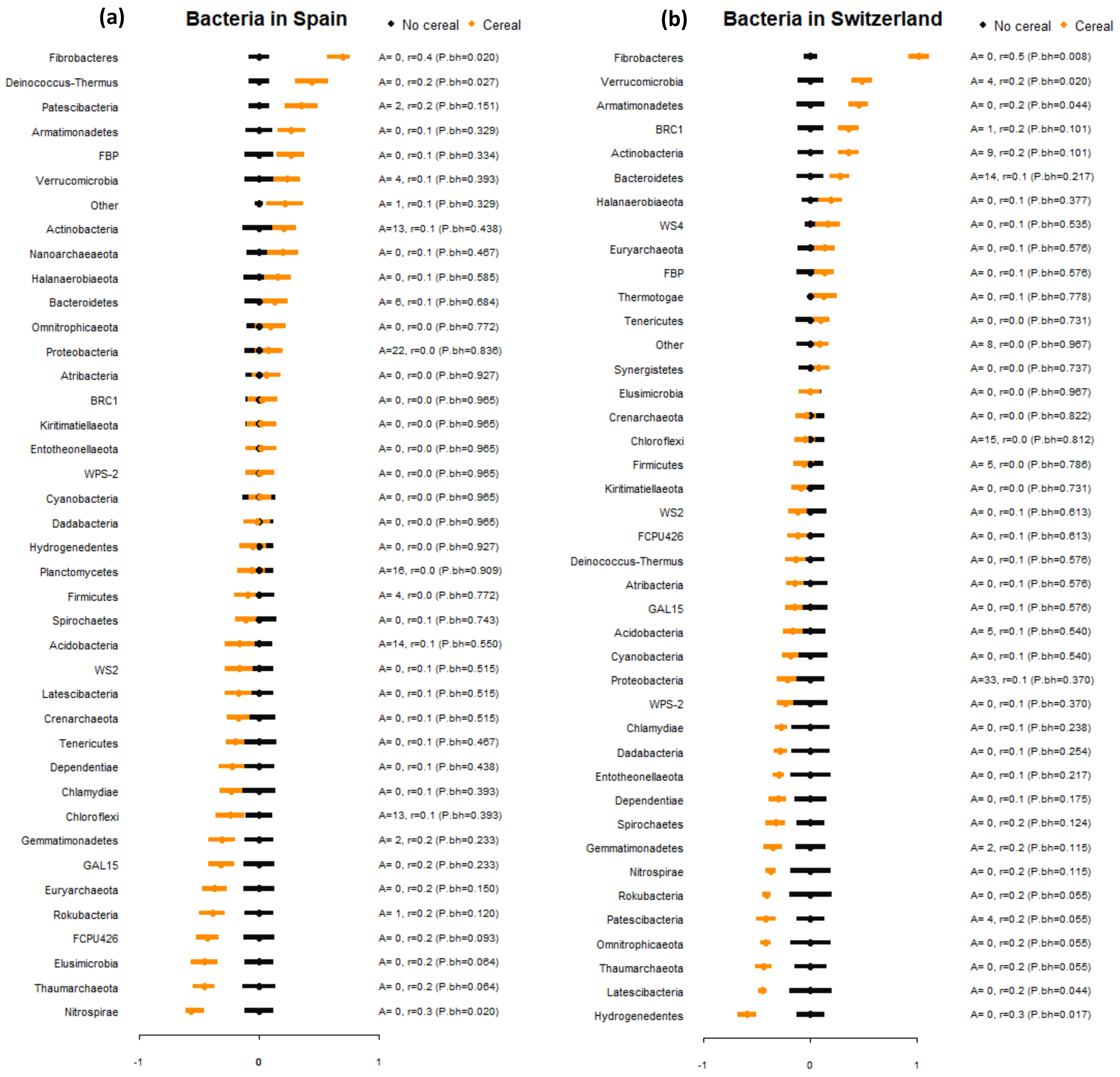

**Figure S10.** Non-randomly distributed taxa along the second (a) and third (b) axes of fungal PCoA decomposition in Switzerland.

CH: Switzerland.

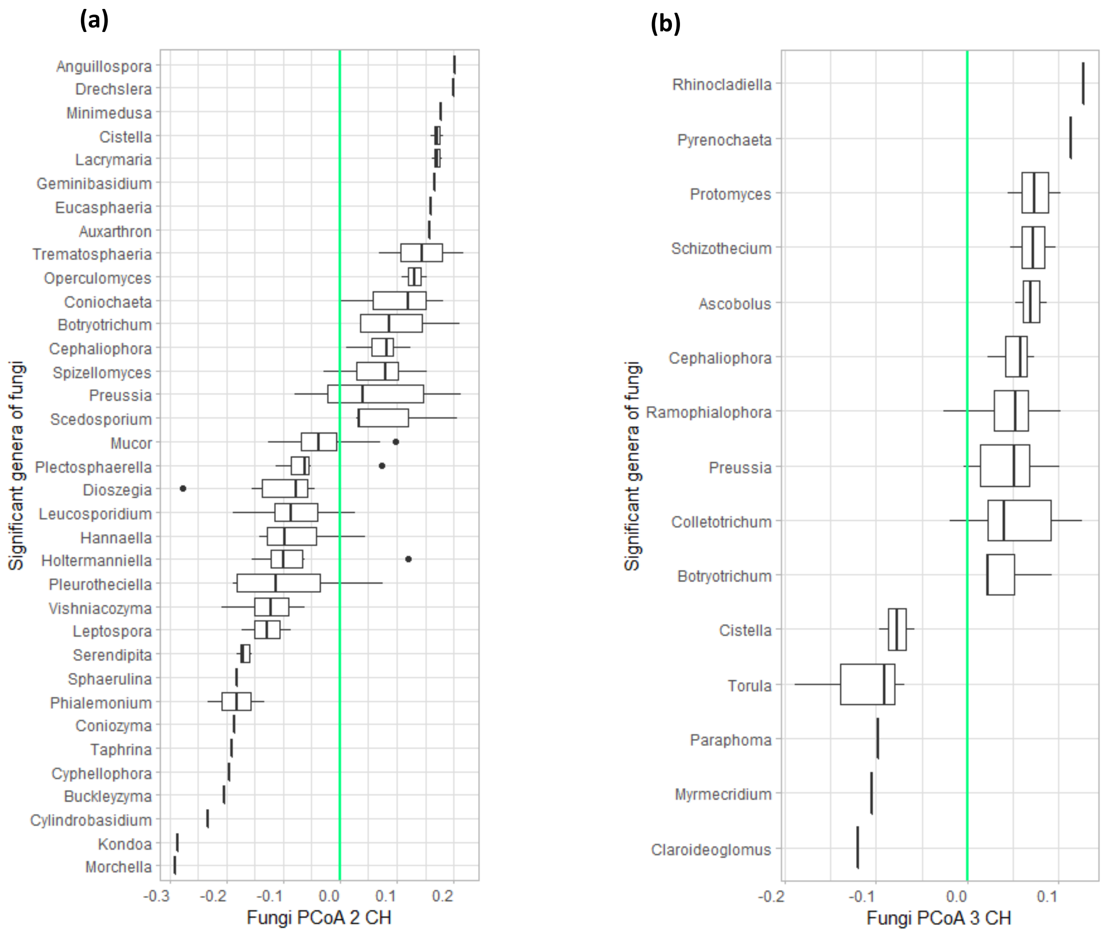

**Figure S11.** Non-randomly distributed taxa along the second axis of bacterial PCoA decomposition in Spain.

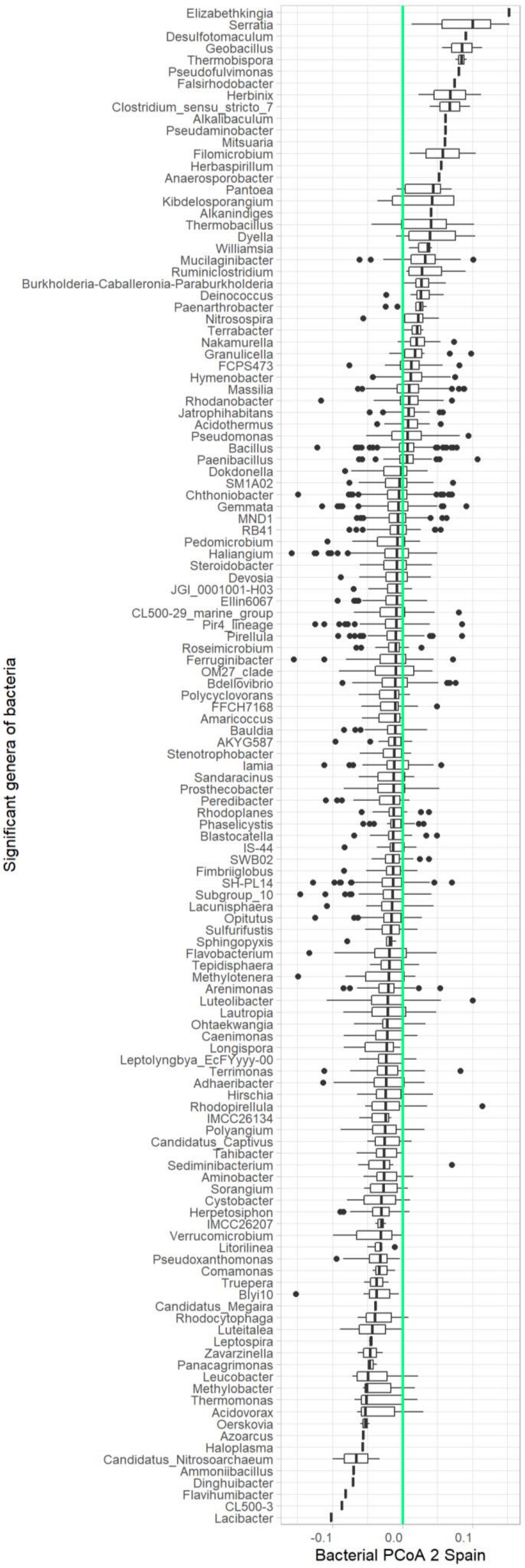

**Figure S12.** Non-randomly distributed taxa along the first (a) and second (b) axes of bacterial PCoA decomposition in Switzerland.

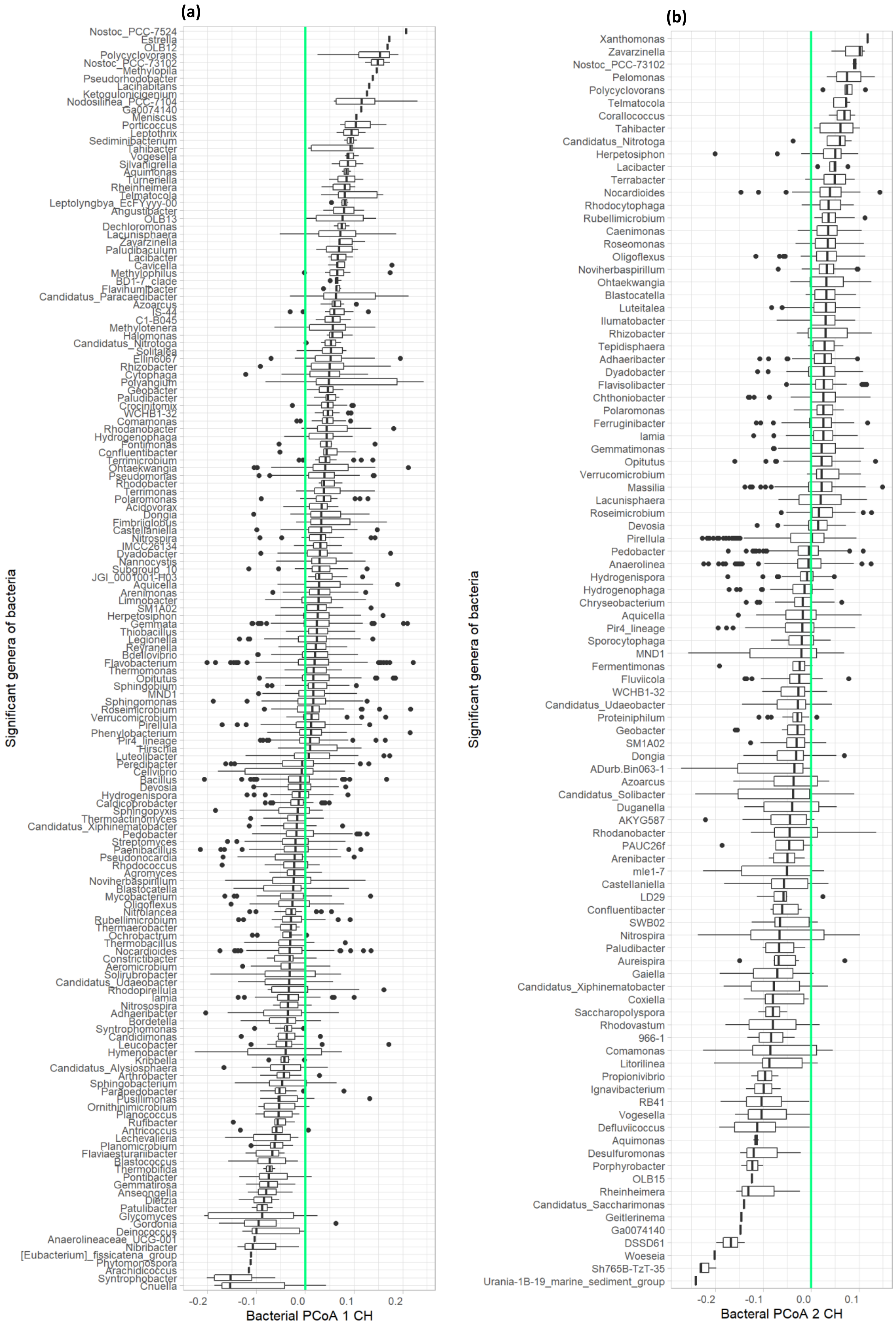
